# Recovery of gene haplotypes from a metagenome

**DOI:** 10.1101/223404

**Authors:** Samuel M. Nicholls, Wayne Aubrey, Arwyn Edwards, Kurt de Grave, Sharon Huws, Leander Schietgat, André Soares, Christopher J. Creevey, Amanda Clare

## Abstract

Elucidation of population-level diversity of microbiomes is a significant step towards a complete understanding of the evolutionary, ecological and functional importance of microbial communities. Characterizing this diversity requires the recovery of the exact DNA sequence (haplotype) of each gene isoform from every individual present in the community. To address this, we present `Hansel` and `Gretel`: a freely-available data structure and algorithm, providing a software package that reconstructs the most likely haplotypes from metagenomes. We demonstrate recovery of haplotypes from short-read Illumina data for a bovine rumen microbiome, and verify our predictions are 100% accurate with long-read PacBio CCS sequencing. We show that `Gretel`’s haplotypes can be analyzed to determine a significant difference in mutation rates between core and accessory gene families in an ovine rumen microbiome. All tools, documentation and data for evaluation are open source and available via our repository: https://github.com/samstudio8/gretel

## Background

Population-level genetic variation enables competitiveness and niche specialization in microbial communities [1]. Recovering the haplotypes of isoforms for a given gene across all organisms in a microbiome would aid evolutionary and ecological insights into microbial ecosystems with biotechnological potential [2, 3, 4]. The concept of haplotyping has been applied to clonal [5] and viral sequence data [6], and the identification and tracking of strains between metagenomic samples [7, 8]. However, there is no method that can recover haplotypes for a particular gene of interest from sequencing data from a microbiome.

Reconstructing population-level variation in microbial communities is limited by difficulties culturing many microbes from the environment [9]. We instead rely on DNA isolated and sequenced directly from an environment (metagenomics). The presence of an unknown number of haplotypes complicates the already computationally difficult (NP-hard) [10] problem of haplotyping [11]. Contemporary haplotyping methodologies are limited to diploid species [12, 13], clonal communities [5, 6] or samples with well-defined genomes [14, 15, 8]. Existing *de novo* analysis pipelines for DNA sequence data generally assume a single individual of origin [16] or, when applied to metagenomic datasets, remove low level variation and produce single consensus sequences which represent an “average” of all the haplotypes present [17]. Even specialized metagenomic assemblers do not aim to solve the problem of haplotype recovery [17, 18, 19]. Naive sequence partitioning approaches such as binning or clustering assembled contigs often reconstruct bins with low quality and genome completeness [20] or require many samples in order to quantify strain abundances [21]. Thus, the problem of recovering haplotypes from a metagenome does not have a formal mathematical definition.

To address this, here we present a formalization of the problem of recovering the set of “gene haplotypes” found across individuals in a mixed microbial population, for any locus of interest. Our formalization defines a computationally tractable problem which we have implemented in `Hansel` and `Gretel`: a Bayesian framework that recovers and ranks gene haplotypes from metagenomic data using evidence of pairs of single nucleotide polymorphisms (SNPs) observed on sequenced reads.

We characterize the performance of our approach on simulated metagenomes, demonstrate its effectiveness on data from a natural microbial community and validate these results using Sanger and Pacific Biosciences sequencing. This empirical evidence demonstrates that the most likely haplotypes recovered from complex metagenomic samples by `Hansel` and `Gretel` represent real previously unknown gene isoforms. We show how this information is neglected by methods that aim to resolve (consensus) genomes from a metagenome, or track strains with marker genes across multiple microbial samples. Finally, to demonstrate that `Gretel` can be used to elucidate valuable biological information, we use `Gretel` to inspect the diversity of haplotypes within a population of *Ruminococcus flavefaciens* in the ovine gut.

## Results

### The terminology for haplotypes in a metagenome can be mathematically defined

Here we provide a mathematical formalization that defines the computational problem of recovering gene haplotypes from metagenomic data with precise language for future work on haplotyping to draw upon. We formally describe a metagenome and offer the *metahaplome* as a term to refer to the set of haplotypes from all organisms for any particular homologous region of interest (such as a gene, or gene cluster) within a metagenomic data set. We define a data structure constructed from pairs of SNPs observed in sequencing reads aligned to a reference or metagenomic assembly and formulate the problem of recovering haplotypes from a metahaplome as a graph traversal problem; whereby traversal of the edges in the graph depend on the observation of SNP pairs. The full mathematical definitions are provided in Supplementary Section 1: succinctly describing the `Hansel` data structure, how it can be used by `Gretel` to construct a graph that permits haplotype recovery, and the equations that transform SNP pair observations into probabilities for selecting which variants to add to a growing haplotype and estimating haplotype quality.

### Hansel and Gretel is suitable for a wide range of short read lengths and read coverage

We evaluated the fidelity of the haplotypes reconstructed by `Gretel` using 6,300 synthetic metahaplomes, representing seven mutation rates in combination with the 3 read sizes and 6 per-haplotype read depths (Figure 1).

**Figure 1:**
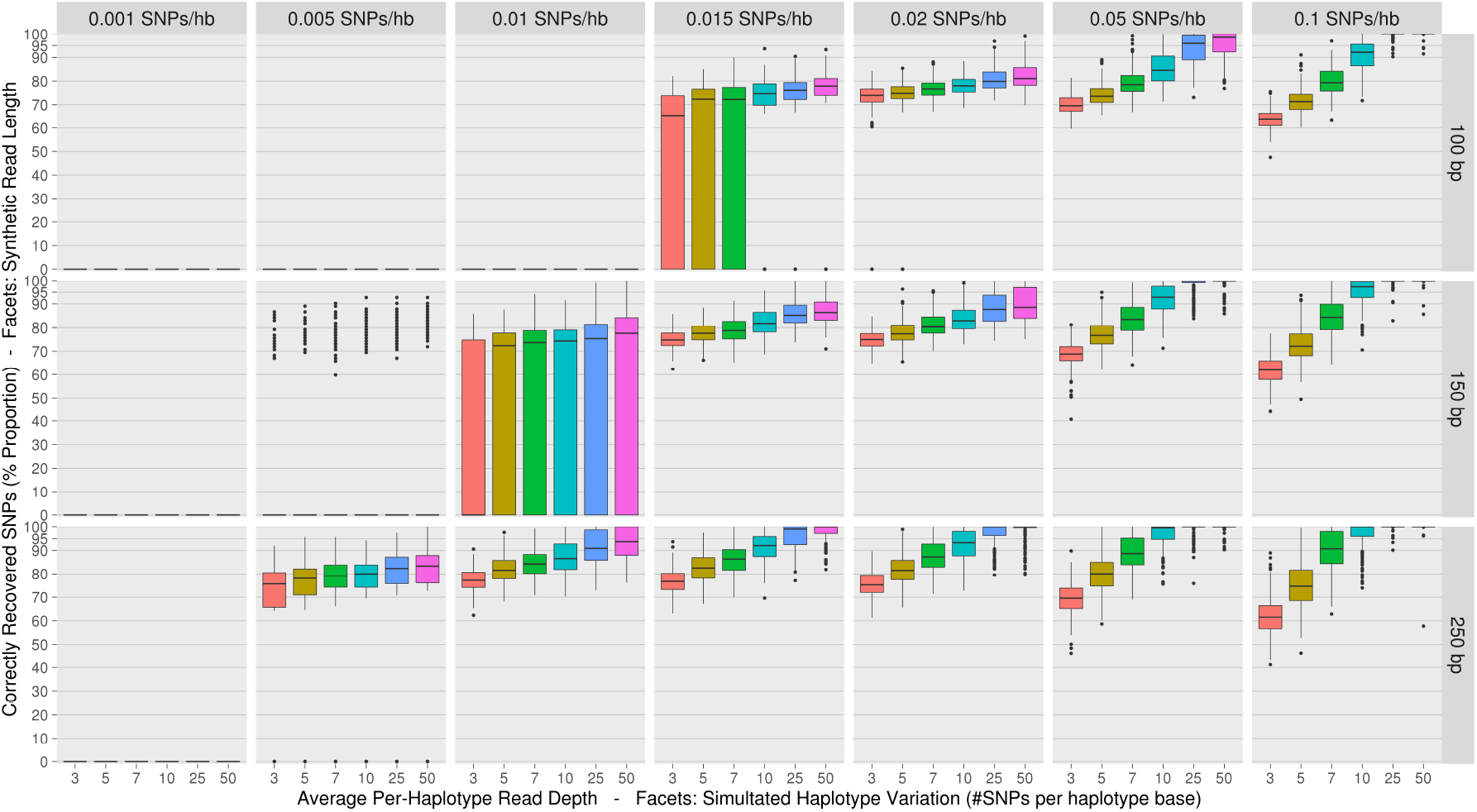
Boxplots summarising the proportion of variants on an input haplotype correctly recovered (y-axes) from groups of synthetic metahaplomes by `Gretel`. Single boxplots present recoveries from a set of five metahaplomes generated with some per-haplotype mutation rate (column facets), over 10 different synthetic read sets with varying read length (row facets) and per-haplotype read depth (colour fill). Each box-with-whiskers summarises the proportion of correctly recovered variants over the 250 best recovered haplotypes (yielded from 50 `Gretel` runs (5 metahaplome replicates *×* 10 read sets), each returning 5 best outputs). We demonstrate better haplotype recoveries can be achieved with longer reads and more dense coverage, as well as the limitations of recovery on data exhibiting fewer SNPs/hb. This figure may be used as a naive lookup table to assess potential recovery rates for one’s own data by estimating the level of variation, with the average read length and per-haplotype depth.

As expected, we found that haplotype recovery improves with longer reads and greater coverage. Unsuccessful recoveries were found to be as a result of at least one pair of adjacent variants failing to be covered by any read, which is a requirement imposed on `Gretel` for recovery of haplotype paths.

We recover perfect haplotypes from data sets with high variation and find that with coverage of *≥* 7x per-haplotype depth, `Gretel`’s haplotypes are more accurate than those in data sets with fewer SNPs (Figure 1).

**Table 1:** `Gretel` haplotypes and their corresponding PacBio circular consensus sequences (CCS). The haplotypes produced by `Gretel` for G31 had no perfectly matching PacBio sequences. The closest match for G31 had 3 mismatches.

| Amplicon | Gretel Haplotype Likelihood | Number of perfect CCS |
| --- | --- | --- |
| G90 | -28.13 | 61 |
| G90 | -50.64 | 16 |
| G123 | -16.65 | 1592 |
| G123 | -27.29 | 458 |
| G123 | -28.60 | 405 |
| G123 | -32.79 | 351 |
| G123 | -36.02 | 845 |

For realistic levels of variation (0.01–0.02 SNPs per haplotype-base [hb]) [22] and per-haplotype read depth of *≥* 7*×*, we can recover haplotypes with a median accuracy of 84.40% (Max = 100%). With at least 0.15 SNPs/hb and a higher per-haplotype coverage of *≥* 25*×*, `Gretel` recovers haplotypes with a median accuracy of 99.85%, with many haplotypes recovered at 100% accuracy.

### Hansel and Gretel is capable of recovering haplotypes from mixed strains

To demonstrate that `Hansel` and `Gretel` can recover haplotypes for strains in a mixed community we used a mock community from Quince *et al.* [7]. The mock community contains 5 *Escherichia coli* strains, and 15 other genomes commonly found in the human gut according to the Human Microbiome Project [23]. The authors made available 16 million synthetic read pairs, generated from the 20 genomes to simulate a typical HiSeq 2500 run [7]. Additionally, Quince *et al.* identified 982 single-copy genes for *E. coli* and provided DNA sequences of all the genes for each of the five *E. coli* strains in their mock community.

Our assembly of the synthetic reads contained 814 of the original 982 single-copy genes. `Gretel` was executed over these 814 genes with the aim of recovering the complete set of 814 gene haplotypes for each of the five *E. coli* strains. `Gretel` recovered full-length haplotypes for the 814 single-copy genes for the 5 strains of *E. coli* with a median accuracy of 99.5% (Figure 2).

**Figure 2:**
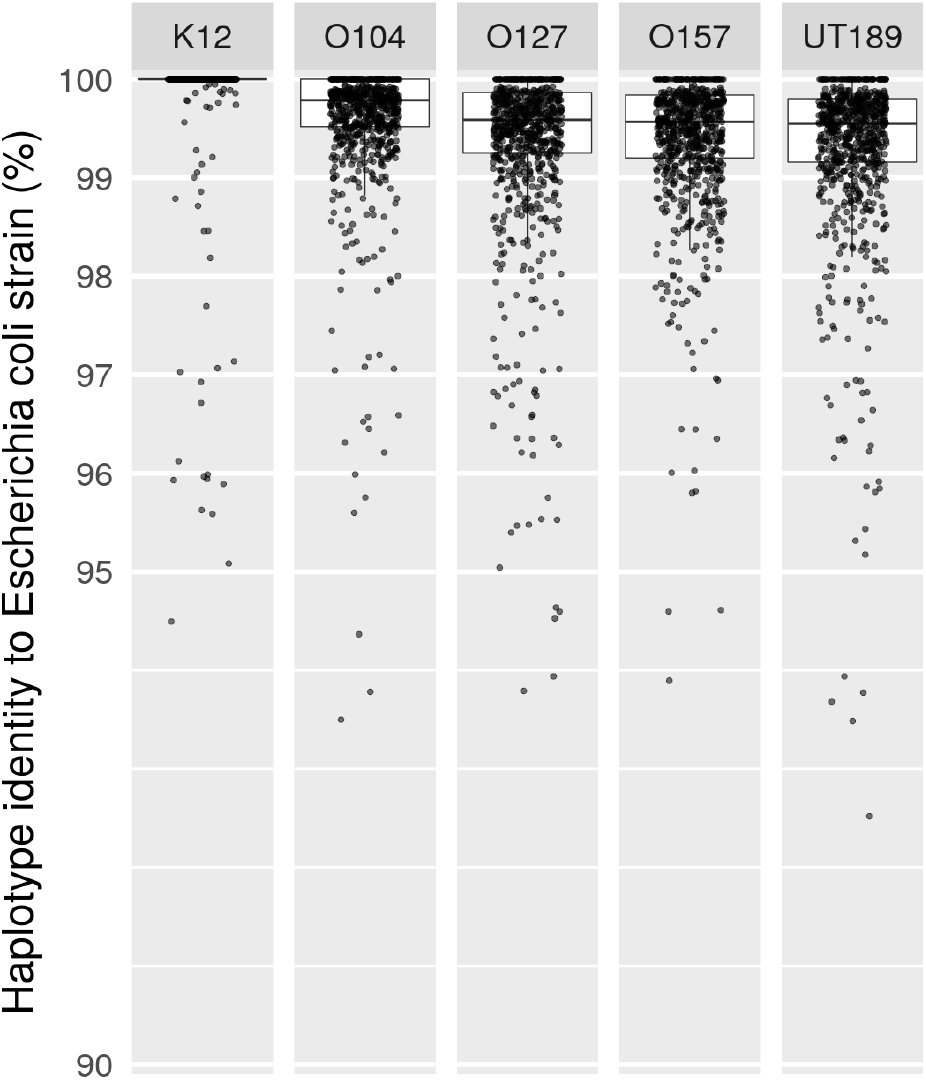
The boxplot summarises the percentage sequence identity (y-axis) of `Gretel` haplotypes recovered from each of the 814 gene sites, to five *E. coli* strains (column facets) known to exist in the mock community. `Gretel` was executed at 814 sites on an assembled mock metagenome, consisting of short-reads generated from five *E. coli* strain haplotypes, and 15 other genomes. The y-axis is truncated at 90%.

### Hansel and Gretel recovers gene haplotypes from a real metagenome

To validate our method empirically, we predicted haplotypes from a natural microbial community sequenced using Illumina short-reads, for three genes of interest (which we refer to as G31, G90 and G123 [Table 2]), and verified their existence by sequencing isolated amplicons with both Sanger and PacBio single-molecule long-read sequencing. PacBio sequencing of the three amplicons yielded 149,003 high-quality circular consensus sequences (CCS).

**Table 2:**
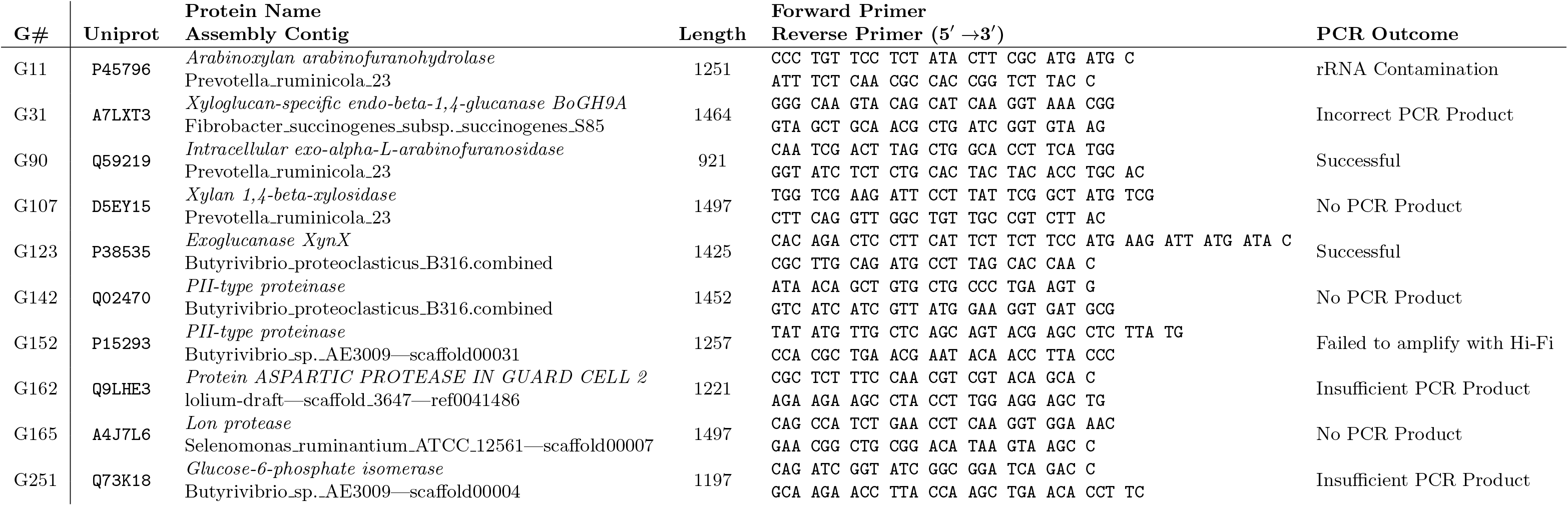
Annotations for the 10 genes that were selected from 259 possible candidates for *in vitro* verification of our `Hansel` and `Gretel` haplotype recovery framework. Three amplicons (G31, G90 and G123) could be generated in sufficient quality and quantity for sequencing.

The CCS reads support `Gretel`’s recovery of five distinct haplotypes for G123, with perfectly matching CCS reads numbering between 351 and 1592. Additionally, the CCS reads support the existence of two G90 haplotypes. The haplotypes with perfect matches to the PacBio CCS reads were those with the top five and top two likelihoods for G123 and G90 respectively, demonstrating the utility of the likelihood scores calculated by `Gretel. BLAST` alignments also verify perfect matches of CCS reads to `Gretel` recovered haplotypes (Supplementary Sections 11.3 and 11.4).

G31 appears to be poorly recovered, with no CCS reads perfectly matching a `Gretel` reconstructed haplotype. Although there exist CCS reads with as few as 3 mismatches against a G31 haplotype from `Gretel`, generally the CCS reads for G31 were highly divergent from any haplotype recovered by `Gretel` (Supplementary Table 3). However, aligning the initial Illumina data to these CCS reads, we observe regions of low coverage and pileup positions which do not agree with the CCS reads, implying that the material that was amplified for PacBio sequencing does not correspond to the short-read Illumina sequencing from which the haplotypes were predicted, despite the amplification originating from the same frozen samples (Supplementary Section 11.6.1). Regardless, it is possible to load and interrogate the `Hansel` matrix that was used by `Gretel` during the run that recovered the G31 haplotypes, and inspect the evidence available for any CCS read. The pairwise SNP information stored by `Hansel` demonstrates that the Illumina reads simply did not provide `Gretel` evidence to recover haplotypes for G31 that would concur with the CCS reads (Supplementary Section 11.6.3). This could potentially arise due to off-target amplification of G31, or a lack of coverage for the G31 loci in the initial Illumina shotgun meta-transcriptome sequencing. This is not a shortcoming of our method: `Gretel` can only recover haplotypes for which there is sufficient sequenced evidence available in `Hansel`.

### Hansel and Gretel reveals the haplotype landscape of a pangenome

To demonstrate the scalability and power of `Hansel` and `Gretel` to generate novel population-level haplotype information, 2 samples of shotgun whole metagenome short-reads from sheep microbiomes were downloaded and aligned to 14 reference sequences for *Ruminococcus flavefaciens*, a common constituent of this gut ecosystem. 29,476,758 reads aligned to the references. The 14 references were annotated with 16,801 CDS regions from which haplotypes could be reconstructed from the metagenomic data. `Gretel` recovered 119,151 haplotypes in total.

The rate of non-synonymous to synonymous mutations (dN/dS) was used to characterize the diversity of the haplotypes recovered for each genes family across the 14 *R. flavefaciens* genomes. Higher dN/dS ratios represent elevated rates of amino-acid substitutions in the `Gretel` generated haplotypes and therefore elevated numbers of protein isoforms in the population, which has been previously linked to niche specificity in the rumen microbiome [1].

When combined with the prevalence of the gene families in the 14 reference genomes this information allows exploration of the relationship of population variation between core and accessory genes for the pangenome of this species in the environments from which the metagenomic samples were taken.

We found that the dN/dS ratio was more variable and statistically higher (Welch Two Sample t-test, one-tailed, *p <* 2.2 *×* 10^*−*16^) for accessory gene families in the pangenome (gene families appearing in 1–3 strains, n=864) than for core gene families (those found in 12–14 genomes, n=324) (Figure 5).

### Hansel and Gretel outperforms clonal/viral haplotyping and strain inference tools on metagenomic data

Application of methods designed for clonal haplotyping to metagenomic data from mixed microbial communities can lead to very poor recovery rates as a clonal population is much simpler than that of a mixed microbial community. To show this, we ran the clonal haplotyping tool `EVORhA` [5] on our Illumina reads for the contig containing the G123 locus. `EVORhA` reconstructed 16 global haplotypes for the contig. Fifteen of these haplotypes were identical for the G123 locus, yielding only two haplotypes differentiated by three SNPs. No PacBio CCS reads supported either of the two `EVORhA` G123 haplotypes. We were unable to run the older PredictHaplo on our own data, a problem also encountered by Pulido-Tamayo *et al.* in their comparison to `EVORhA`.

A more recent approach for the related problem of viral quasi-species recovery has been implemented in `SAVAGE` [6] and its successors `Virus-VG` [24] and `VG-Flow` [25]. Baaijens *et al.* claimed performance improvements for `SAVAGE` over both `PredictHaplo` [26] and `ShoRAH` [27] using a benchmark lab mixture of five strains of HIV. To compare we evaluated `Gretel` on the same HIV data. Our results are presented in full in Supplementary Section 13, but in summary, we found that `Gretel` makes perfect recoveries from this sequenced laboratory mix of five strains (Supplementary Table 7).

Alternative strategies for metagenomes focus on strain inference. Given a database of genes known to occur in single-copy for a species, `DESMAN` identifies SNP frequency patterns over multiple samples (*>*50 samples preferred) to identify and compare the presence of strains between samples. The use of frequencies observed across samples means restricting analyses to single copy genes, as multiple copies would distort the frequencies; limiting its use to identifying marker genes only. The necessity for many samples and the restriction of strain inference on single-copy genes meant we were unable to run `DESMAN` on our own testing data, as `Gretel` is not restricted to single-copy genes for one species, and our data represents the scenario of a single microbial sample, rather than a large set of samples expected by `DESMAN`. In contrast, `Gretel` was able to make excellent recoveries on the mock community data generated by Quince *et al.* for the `DESMAN` paper (Figure 2).

## Discussion

Here we present `Hansel`, a data structure designed to efficiently store variation observed across ordered sequenced reads, and `Gretel`, a Bayesian algorithm that leverages `Hansel` for the recovery of haplotypes from a metagenome.

`Gretel` is the only tool capable of recovering haplotypes from metagenomes. `Gretel` requires no configuration, has no parameters, requires no pre-processing of reads, does not discard observed information and is designed for data sets where the number of haplotypes is unknown. Importantly, it also addresses the limitations that make existing haplotyping methods unsuitable for metagenomic analysis:

- assuming that the solution is a pair of haplotypes from diploid parents, and discard/alter observations until a pair of haplotypes can be determined [16]
- discarding SNP sites that feature three or more alleles as errors [28]
- generating an unrealistically large number of unranked potential haplotypes [4, 29]
- are too computationally expensive for high-depth short read data sets [30]
- require a good quality reference genome [31]
- are no longer maintained/are specific to certain data/cannot be installed [26]

Through our initial simulations we found that contrary to the expectation that high levels of variation would make the recovery of haplotypes more challenging (as the potential haplotype search space is larger), an abundance of variation actually provides more information for the likelihood approach implemented in `Gretel`, making it suitable for the analysis of complex microbiomes.

### Hansel and Gretel works with real sequencing data

`Gretel` easily tackles the problem of recovering haplotypes for the five different *E coli.* strains over 814 loci in the Quince *et al.* mock community, from data emulating a real sequencing run. In comparison, the binning step of the `DESMAN` pipeline on the same mock community data led to a majority of the single-copy genes being discarded from the assembly, leaving only 372 (of 982) genes for their own downstream strain inference analysis [7]. `Gretel` does not rely on pre-processing and binning. Additionally, we are not limited to the study of single-copy gene families which represent as few as only 1% of all gene families in microbial genomes [32]. For the typical use-case where one lacks sufficient reference sequences or sufficient knowledge of the taxa in a microbial environment, `DESMAN` falls back to the use of 36 single-copy core genes that are universal to all species, severely limiting its utility to the study of DNA sequences found in a microbiome.

Our analysis of the HIV laboratory mix shows the applicability of `Gretel` to real sequencing reads and the benefit of its haplotype likelihood scores. `SAVAGE` is the most accurate tool of the three published by Baaijens *et al.* that have been used to produce quasi-viral haplotypes from the same HIV laboratory mix standard. `SAVAGE` implements an overlap assembly approach that naively creates haplotypes for every possible branching point in the aligned reads, without considering long range information or determining the likelihood of the haplotypes being generated. Even though both `Gretel` and `SAVAGE` return many haplotypes (`Gretel`: 226 vs. `SAVAGE`: 846), `Gretel`’s likelihood approach allows its haplotypes to be ranked and those with the worst likelihoods can be discarded (Supplementary Figure 9).

Our PacBio sequencing generated 61303 distinct CCS sequences, suggesting that not even highquality DNA sequencing can be used to haplotype alone. We observe CCS reads aligned to our G123 and G90 haplotypes with several mismatches (Supplementary Table 2), but can only speculate as to whether these alternative sequences have arisen as a result of error during reverse transcription which became amplified during PCR, an error during the PCR process itself, PacBio sequencing error or are real mutations from a true haplotype. Such true haplotypes could have been missed by `Gretel`, or were not sequenced to an appropriate depth in the original Illumina sample to have enabled recovery. Our *in vitro* analysis of three amplicons recovered from the bovine rumen microbiome validates that `Gretel` perfectly recovers haplotypes, and that the haplotypes with the best likelihoods are the haplotypes best supported by our validation sequencing data.

### Hansel and Gretel steps beyond strain inference and recovers haplotypes

Species and strain identification is a common theme in recent tools developed to analyze metagenomic data and is sometimes confused with haplotyping. Such tools use specific sets of single-copy marker genes to estimate abundances [7, 8, 33]. Other methods map reads or their component k-mers to pre-built databases of specific species or strains of interest [34, 35]. These tools are not designed to solve the problem of recovering all haplotypes from all genes in a mixed microbial community.

For example, `ConStrains` [8] is a popular state-of-the-art tool that aims to identify strain profiles based on variation observed over core gene markers, and reconstruct their phylogeny, within a metagenomic sample. These profiles act as a barcode that can be used to identify the presence of a strain across a series of samples, and permit the construction of a strain phylogeny, however no haplotypes are generated.

Strain inference methods like `Constrains` and `DESMAN` aim to infer gene counts across the strains within a pangenome, but do not resolve haplotypes. Our pangenome analysis shows that with `Gretel`’s ability to recover haplotypes, we can now go beyond counting the size of the core and accessory genomes and actually calculate mutation rates at the nucleotide level. `Gretel` can provide a landscape overview of the diversity present across every gene in a pangenome.

### Hansel and Gretel is a tractable method for haplotype recovery

Our framework has been designed for the recovery of haplotypes from a loci of interest in a metagenome (such as a gene), but with sufficient observed variation (*i.e.* each adjacent SNP pair is bridged by at least one read) and coverage, our approach could reconstruct haplotypes for significantly longer regions if desired.

The storage of SNP pair observations in the `Hansel` data structure allows for effective compression of the sequenced reads. Once the matrix has been populated, the reads are no longer required. The matrix can also be saved to disk for future interrogation. `Gretel` employs a probabilistic algorithm that is designed with `Hansel`’s structure in mind, leveraging pairs of co-occurring SNPs to build chains of SNPs that form haplotypes. The algorithm is purposely formulated to employ the benefit of Naive Bayes, which significantly reduces the number of calculations required to select the next SNP in the haplotype recovery process, improving performance.

Ultimately, haplotype recovery is bound by the quality of the data available: both the choice of reference and read alignment will exert influence over how many and how accurately haplotypes in a given metahaplome can be recovered. As stated by Lancia, even without error, there are scenarios where data is insufficient to successfully recover haplotypes and the problem is rendered impossible [11].

Regarding time and resource requirements, `Gretel` is designed to work on all reads from a metagenome that align to some region of interest on the metagenomic assembly. Typically these subsets are small (on the order of 10-100K reads) and so our framework can recover haplotypes on an ordinary desktop in minutes, without significant demands on disk, memory or CPU. Run-times on data with very deep coverage (*>* 5000*×*), or many thousands of SNPs, such as the HIV laboratory mix, run in the order of hours, but can still be executed on an ordinary desktop computer.

## Conclusion

The microbial communities with which we share our world and bodies have diverse functions that are widely shared between bacterial strains and species via horizontal gene transfer. Evidence appears to indicate that in complex microbiomes, acquiring a single copy of a gene for a competitive function may not be enough to confer an advantage alone, but rather it is the breadth of functional variants of the gene at the population-level that drives competitiveness. This gene-level population variation is not captured by the assembly of consensus reference sequences or species-specific pangenomes, supporting the need for a more gene-centric view of the microbiome.

`Gretel` addresses this need, offering a gene-centric method for recovering haplotypes from a metagenome with far reaching use cases, including directed enzyme engineering, niche and competition analysis, pan-genome landscape analysis, assembly polishing, analysis of horizontal gene transfer and design of minimal communities.

## Methods

### The metahaplome

We have provided a detailed mathematical definition of the metahaplome in Supplementary Section 1. To enable recovery of a metahaplome from a metagenomic sample we require:

- *g*, a known **target** DNA sequence of interest
- *c*[*i*: *j*], a **region** of contig *c*, identified as having sufficient similarity to *g* by ∆(*c, g*)
- *A*_*c*[*i*:*j*]_, the **alignments** of the set of reads *R* against the contig region *c*[*i*: *j*]
- *S*_*c*[*i*:*j*]_, the genomic positions determined to be **variants** over the region *c*[*i*: *j*]

A set of contigs *C* are generated by assembling sequenced reads with an assembler such as `flye` [36]. Using ∆, one may locate a gene of interest *g*, on a contig *c ∈ C* by similarity search or gene prediction (*e.g.* `prokka`). We refer to gene *g* as the *target*. We want to recover the most likely haplotypes of *g* that exist in the metahaplome Γ_*g*_, using the observations of co-occurring variants observed across the reads that align to *c*[*i*: *j*].

A subset of reads that align to the target region can be determined using a short read alignment tool such as `bowtie2` [37]. Reads that fall outside the region of interest (*i.e.* reads that do not cover any of the genomic positions on *c* that are associated to the target) can be safely discarded: they do not provide relevant evidence for the recovery of haplotypes on the region of interest.

Variation at single nucleotide positions across reads along the target, can then be called with a SNP calling algorithm such as that provided by `samtools` [38] or `GATK` [39]. However, to avoid loss of information arising from the diploid bias of the majority of SNP callers [28], our methodology aggressively considers any heterogeneous site as a SNP using `gretel-snpper`.

The combination of aligned reads, and the locations of single nucleotide variation on those reads can be exploited to recover real haplotypes in the metagenome: the **metahaplome**.

### Hansel: A novel data structure

We present **Hansel**, a probabilistically-weighted, graph-inspired, novel data structure. `Hansel` is designed to store the number of observed occurrences of a symbol *α* appearing at some position in space or time *i*, co-occurring with another symbol *β* at another position in space or time *j*. For our approach, we use `Hansel` to store the number of times a SNP *α* at the *i*’th variant of some contig *c*, is observed to co-occur (appear on the same read) with a SNP *β* at the *j*’th variant of the same contig. `Hansel` is a four dimensional matrix whose individual elements *H*[*α, β, i, j*] record the number of observations of a co-occurring pair of symbols (*α_i_, β_j_*).

### The Hansel matrix is different from the typical Lancia SNP matrix

Our representation differs from the typical SNP matrix model [11] that forms the basis of many of the surveyed approaches. Rather than a matrix of columns representing SNPs and rows representing reads, we discard the concept of a read entirely and aggregate the evidence seen across all reads by genomic position.

At first this structure may appear limited, but the data in *H* can easily be exploited to build other structures. Consider *H*[*α, β*, 1, 2] for all symbol pairs (*α, β*). One may enumerate the available transitions from space or time point 1 to point 2. Extending this to consider *H*[*α, β, i, i* + 1] for all (*α, β*) over *i*, one can construct a simple graph *G* of possible transitions between all symbols. In our setting, *G* could represent a graph of transitions observed between SNPs on a genomic sequence, across all reads. Figure 3 shows how the `Hansel` structure records information about SNP pairs, and shows a simple graph constructed from this information.

**Figure 3:**
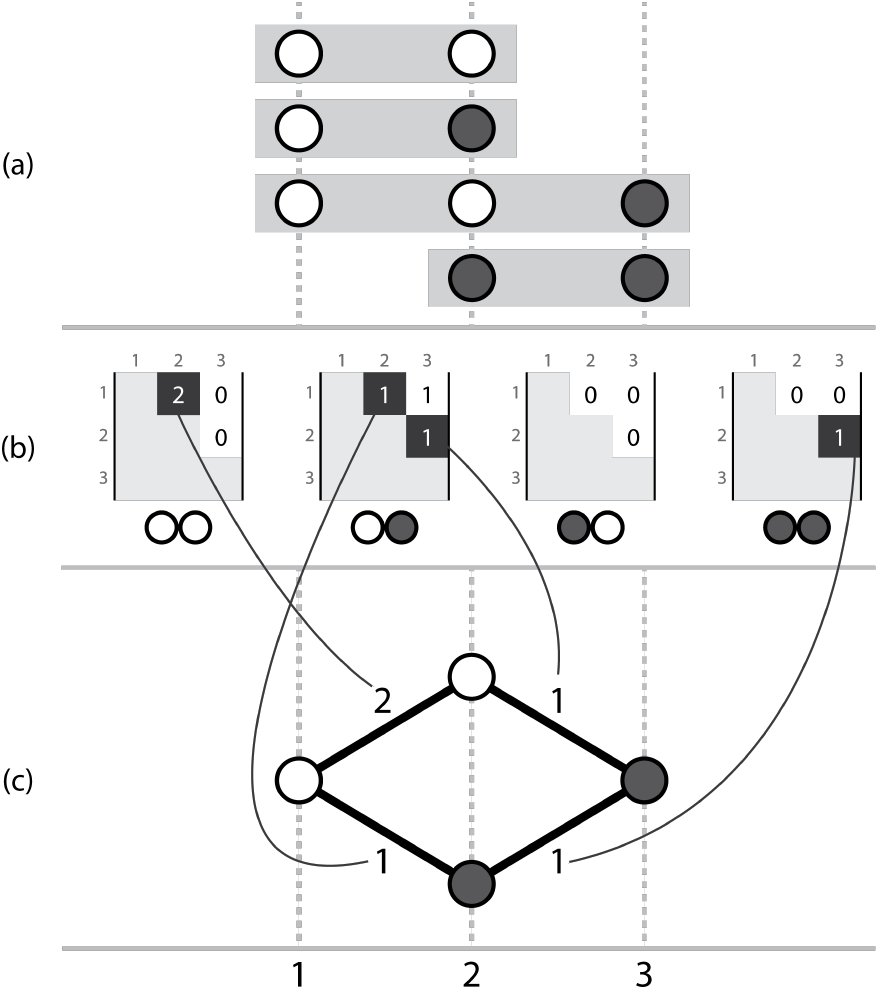
Three corresponding representations, (a) a set of aligned short read sequences, with called variants, (b) the actual `Hansel` structure where each possible pair of symbols (00, 01, 10, 11) has a matrix storing counts of occurrences of that ordered symbol pair between two genomic positions across all of the aligned reads, (c) a simple graph that can be constructed by considering the evidence provided by adjacent variants. Note this representation ignores evidence from non-adjacent pairs, which is overcome by the dynamic edge weighting of the `Hansel` data structure’s interface.

Intuitively, one may traverse a path through *G* by selecting edges with the highest weight in order to recover a series of symbols that represent an ordered sequence of SNPs that constitute a haplotype in the metahaplome. The weight of an edge between two nodes may be defined as the number of reads that provide direct evidence for that pair of SNP values occurring together.

### The Hansel structure allows us to consider information from non-adjacent SNPs

Although the analogy to a graph helps us to consider paths through the structure, the available data cannot be fully represented with a graph such as that seen in Figure 3 alone. A graph representation defines a constraint that only considers pairs of adjacent positions (*i, i* + 1) over *i*. Edges can only be drawn between adjacent SNPs and their weightings cannot consider the evidence available in *H* between non-adjacent SNP symbols. Without considering information about non-adjacent SNPs, one can traverse *G* to create paths that do not exist in the observed data set, as shown in Figure 4. To prevent construction of such invalid paths and recover genuine paths more accurately, one should consider evidence observed between non-adjacent symbols when determining which edge to traverse next.

**Figure 4:**
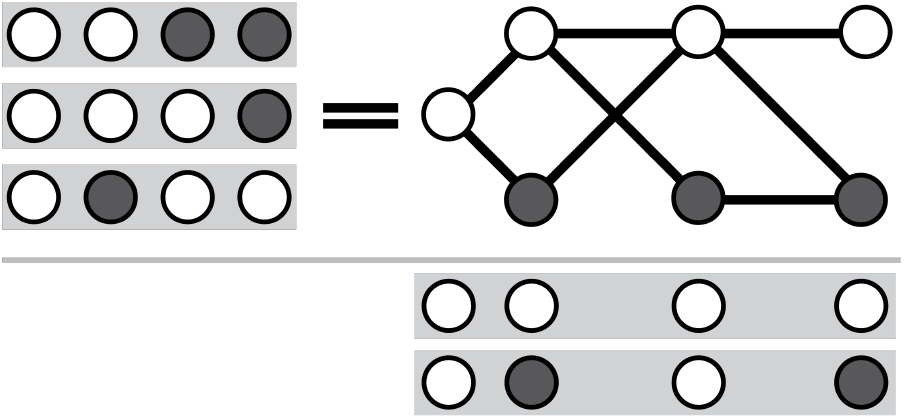
Considering only adjacent SNPs, one may create paths for which there was no actual observed evidence. Here, the reads *{*0011, 0001, 0100*}* do not support either of the results *{*0000, 0101*}*, but both are valid paths through a graph that permits edges between pairs of adjacent SNPs.

### Edge weights in the Hansel graph are dependent on the current path

The `Hansel` structure is designed to store pairwise co-occurrences of all SNPs (not just those that are adjacent), across all reads. We may take advantage of the additional information available in *H* and build upon the graph *G*. Incorporating evidence of non-adjacent SNPs in the formula for edge weights allows decisions during traversal to consider previously visited nodes, as well as the current one.

That is, given a node *i*, the decision to move to a symbol at *i* + 1 can be informed not only by observations in the reads covering positions (*i*, *i* + 1), but also (*i −* 1, *i* + 1), (*i −* 2, *i* + 1), and so on. Such a scheme allows for the efficient storage of some of the most pertinent information from the reads, and allows edge weights to dynamically change in response to the path as it has been constructed thus far. Outward edges between (*i*, *i* + 1) that would lead to the construction of a path that does not exist in the data can now be influenced by observations in the reads beyond that of the current node and the next. Our method mitigates the risk of constructing paths which do not truly exist.

The consideration and storage of pairwise SNPs fits well with the Naive Bayes model employed to simplify the potentially expensive calculation of conditional probabilities (Supplementary Section 4).

Although we describe `Hansel` as “graph-inspired”, allowing edge weights to depend on the current path through *G* itself leads to several differences between the `Hansel` structure and a weighted directed acyclic graph. Whilst these differences are not necessarily disadvantageous, they do change what we can infer about the structure. The structure of the graph is effectively unknown in advance. That is, not only are the weights of the edges not known ahead of traversal (as they depend on that traversal), but the entire layout of nodes and edges is also unknown until the graph is explored (although, arguably this would be true of very large simple graphs too). Indeed, this means it is also unknown whether or not the graph can even be successfully traversed.

Also of note is the fact that the graph is dynamically weighted. The current path represents a memory that affects the availability and weights of outgoing edges at each node. Edge weights are calculated probabilistically *during* traversal. They depend on the observation of SNP pairs between some number of the already selected nodes in the path, and any potential next node. Supplementary Section 3 provides the equation and intuition for the probabilistic calculation of edge weights.

In exchange for these minor caveats, we have a data structure that permits graph-like traversal that is intrinsic to our problem definition, whilst utilising informative pairwise SNP information collected from observations on raw metagenomic reads. `Hansel` fuses the advantages of a graph’s simple representation (and its inherent traversability) with the ability to efficiently store pertinent information by considering only pairs of SNPs across all reads.

**Figure 5:**
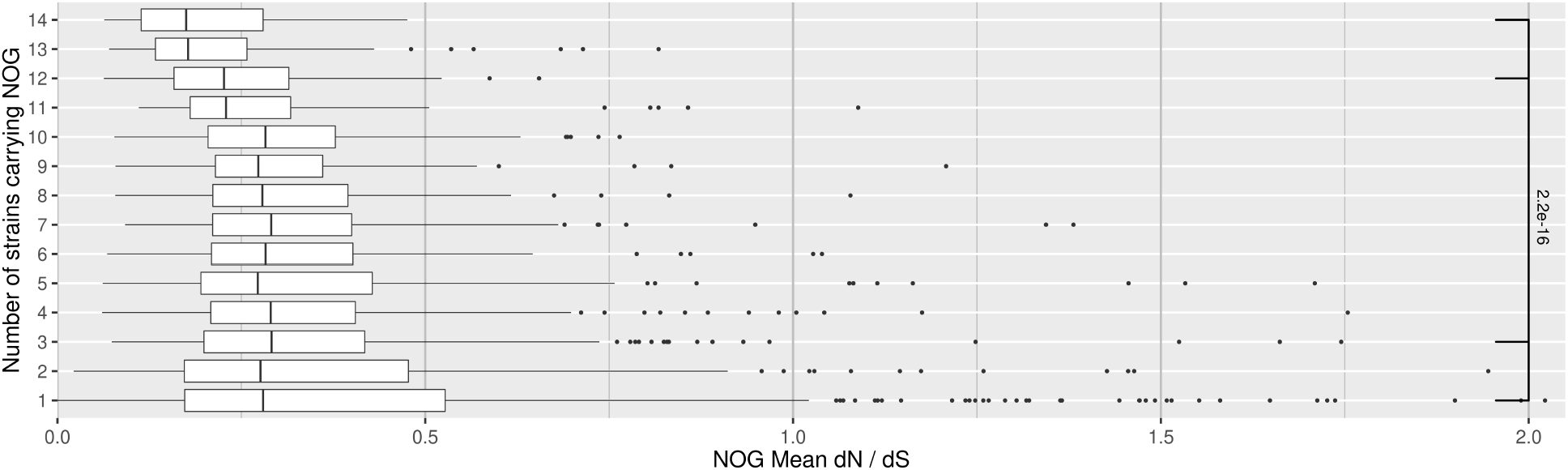
Boxplot describing the distribution of dN/dS ratios for EggNOG gene families that appear on a given number of *R. flavefaciens* references. Each box-and-whiskers shows the average dN/dS ratio for all haplotypes emitted from regions assigned by emapper to a particular gene family (x-axis), that appears on a given number of strain references for *R. flavefaciens* (y-axis). We can the estimate core and accessory genes families by counting those that appear on fewer strains as accessory, and conversely those that appear on more strains as core. We observe a statistically significant difference between average dN/dS ratio between gene families on three or fewer strains and gene families on twelve or more strains.

### Gretel: An algorithm for recovering haplotypes from metagenomes

We introduce `Gretel`, an algorithm designed to interface with the `Hansel` data structure to recover the most likely haplotypes from a metahaplome. To obtain likely haplotypes, `Gretel` traverses the probabilistic graph structure provided by `Hansel`, selecting the most likely SNPs at each possible node (*i.e.* traversing edges with the greatest probability), given some subset of the most recently selected nodes in the path so far. At each node, an *L*’th order Markov chain model is employed to predict which of the possible variants for the next SNP is most likely, given the last *L* variants in the current path. Execution of `Gretel` can be broken into the following steps:

1. Parse the read alignments and retain only the bases that cover SNP sites, discarding any conserved base positions as they provide no haplotype information.
2. Populate the `Hansel` structure with all pairwise observations from each of the reads.
3. Exploit the `Hansel` graph API to incrementally recover a path until a variant has been selected at each SNP position:

- Query for the available transitions from the current position in the graph to the next SNP
- Calculate the probabilities of each of the potential next variants appearing in the path given the last *L* variants
- Append the most likely variant to the path and traverse the edge
4. Report this path as a haplotype and then remove the information for this path from the data by reweighting observations that contributed to this path. This will allow for new paths to be retrieved next.
5. Repeat (3-4) until the graph can no longer be traversed or an optional additional stopping criterion has been reached.

### Construction of haplotype paths

Haplotypes are reconstructed as a path through the `Hansel` structure, one SNP at a time, linearly, from the beginning of the sequence. At each SNP position, the `Hansel` structure is queried for the variants that were observed on the raw reads at the next position. `Hansel` also calculates the conditional probabilities of each of those variants appearing as the next SNP in the sequence, using a Markov chain of order L that makes its predictions given the current state of the observations in the `Hansel` matrix and the last *L* selected SNPs. `Gretel`’s approach is greedy: we only consider the probabilities of the next variant. Our razor is to assume that the best haplotypes are those that can be constructed by selecting the most likely edges at every opportunity.

### Gretel re-weights the Hansel graph to find multiple haplotypes

Whilst our framework is probabilistic, it is not stochastic. Given the same `Hansel` structure and operating parameters, `Gretel` will behave deterministically and return the same set of haplotypes every time. However, we are interested in recovering the metahaplome of multiple, real haplotypes from the set of reads, not just one haplotype. `Hansel` exposes a function in its interface for the reweighting of observations. Once a path through the graph is completed (a variant has been chosen for all SNP sites), the observations in the `Hansel` matrix are reweighted by `Gretel`.

Currently, `Gretel` reduces the weight of each pairwise observation that forms a component of a completed path - in an attempt to reduce evidence for that haplotype existing in the metahaplome at all, allowing evidence for other haplotypes to now direct the probabilistic search strategy.

### Description of Gretel’s output files

`Gretel` outputs recovered sequences as FASTA, requiring no special parsing of results to be able to conduct further analyses. In addition to the sequences themselves, `Gretel` outputs a ‘crumbs’ file, which contains metadata for each of the recovered sequences: log probability of that sequence existing given the reads, how much of the evidence in `Hansel` the sequence was supported by, and how much of the evidence was reweighted as a result of that path being chosen.

Currently, `Gretel` will recover paths out of the remaining evidence until it encounters a node from which there is no evidence that can inform the next decision.

### Hansel and Gretel is suitable for a wide range of short read lengths and read coverage

To first test our approach on data with well-defined and controllable read properties, but still posing a recovery problem, `seq-gen` [40] was used to generate sets of DNA sequences that would serve as haplotypes of a synthetic metahaplome.

`seq-gen` simulates the evolution of a nucleotide sequence along a given phylogeny. For testing `Gretel`, we provided a star shaped guide tree with uniform branch lengths, such that all haplotypes would be equally dissimilar to each other. These uniform branch lengths correspond to the rate of per haplotype base (hb) nucleotide heterogeneity. Thus, each taxa in the guide tree will have a DNA sequence based upon the evolution of the given starting sequence, following simulated evolution at the given rate.

The same starting sequence was shared by all of our generated trees. We used a randomly generated sequence of 3000 nt with 50% GC content. We fixed the number of taxa in the trees at five, but varied the mutation rate across seven levels. 35 trees were generated (7 mutation rates and 5 replicates), each containing five sequences mutated at the same rate, from the original 3000 nt sequence. Each of the resulting 35 sets of five mutated DNA sequences represent a metahaplome from which the five haplotypes must be recovered by `Gretel`.

We generated synthetic reads from each of the five sequences in the metahaplome, varying both the read length and per-haplotype read depth (*i.e.* the average coverage of each haplotype). For each read length and coverage parameter pair, ten read sets were generated, to amortise any effect on haplotype recovery introduced by the alignments of the reads themselves. We generated 6300 read sets (3 read sizes, 6 per-haplotype depth levels, 7 mutation rates, 10 read replicates, 5 tree replicates). Synthetic reads and their corresponding alignments to the starting sequence reference, are generated *in silico* with our Python tool: (`shredder`).

As pileups of the generated reads typically feature many tri- or tetra-allelic sites (especially as mutation rate increases), most SNP-calling tools would be inappropriate for our task. To avoid diploid tool bias, we use `gretel-snpper` to aggressively call for variants in the generated SAM, outputting a VCF record for any heterogeneous site.

To evaluate the accuracy of a run of `Gretel`, each known input haplotype is compared pairwise to each of the recovered output haplotypes. Each input haplotype is matched to a corresponding “best” recovered haplotype. Best is defined as the output haplotype that yields the smallest Hamming distance from a given input haplotype. For each synthetic metahaplome, we perform a multiple sequence alignment with `MUSCLE` [41] to determine the definitive SNP positions. When calculating Hamming distance, we consider only these corresponding positions. That is, we exclude the comparison of homogeneous sites from the evaluation metric, to ensure we only consider our accuracy on positions that required recovery. For our results we report the proportion of SNPs that were correctly recovered by `Gretel`, expressed as a percentage.

Comparing sites enumerated by the multiple sequence alignment of the original haplotypes, as opposed to the VCF of each individual read set ensures `Gretel` is penalised when a SNP has not been called from the read set.

Regardless of quality, all input haplotypes are assigned a best output haplotype. An output haplotype may be the best haplotype for more than one input. If more than one output haplotype has the same Hamming distance, the first that was found is chosen. If `Gretel` could not complete at least one haplotype (*i.e.* a pair of adjacent SNP positions were not covered by at least one read), all input haplotypes are awarded 0%.

### Metahaplomes from a mock community

In lieu of a true, annotated metagenome, we sourced a benchmark microbial community from Quince *et al.* [7]. The community consists of 5 *Escherichia coli* strains, and 15 other genomes commonly found in the human gut according to the Human Microbiome Project (HMP). The community is defined in the supplement to the author’s original manuscript. In their work, 1.504 *×* 10^9^ reads were generated from the 20 genomes, distributed across 64 paired-end samples (11.75 million read pairs per sample). The `ART` read simulator was configured to simulate a “typical *HiSeq* 2500 run”. As part of their preprint, the authors made available a subset of the generated mock community. The subset contains 16 samples, with 1 million read pairs each, for a total of 32 million reads. Reads were assembled with MEGAHIT [42], using default parameters, as per the author’s recommendations (github.com/chrisquince/DESMAN/blob/master/complete_example/README.md, commit `9045fe2`). Following the example, we discarded assembled contigs shorter than 1 kbp, to yield an assembly described by Supplementary Table 1.

The original Quince *et al.* paper also identified 982 single-copy genes for *E. coli*. Additionally the work provided DNA sequences for all 982 genes, for each of the five different *E. coli* strains found in the mock community. Single-copy genes were mapped to the metagenomic assembly with blastn, with alignments requiring a threshold of at least 75% of the average length of the five haplotypes for each gene. We limited our analysis to the 814 loci for which all five strains could be aligned against the metagenomic assembly, to ensure that `Gretel` would have the opportunity to recover them.

Reads across the 16 samples were concatenated to create one paired-end sample containing 16 million read pairs. The reads were then also mapped to the metagenomic assembly with `bowtie2` (`--sensitive-local`).

`Gretel` was then executed on the aligned reads, once for each of the 814 identified single-copy gene loci with the aim of recovering the five strain haplotypes from the synthetic short-reads. SNPs were called over each region using the `gretel-snpper` method previously described. Performance was measured with a blastn alignment between the known five strain haplotypes, and the `Gretel` recovered haplotypes. In the same fashion as our synthetic evaluation, each input haplotype is assigned a best output haplotype, and an output haplotype may be the best haplotype for more than one input. For each strain, we report the sequence identity of the best haplotype for each of the 814 loci.

### Recovery from a real microbiome

A previous experiment [43] isolated RNA from 32 rumen samples from 3 cows over 6 timepoints (0, 1, 2, 4, 6 and 8 hours) after feeding. In preparation for metatranscriptomic sequencing, the polyA fraction was removed (MicroPoly(A)Purist, Ambion). 18S rRNA was also removed (both RiboMinus Plant Kit and Eukaryote Kit, Invitrogen). 16S rRNA was removed (Ribo-Zero rRNA removal kit (bacteria), Epicentre) all according to the manufacturer’s protocols. The resulting enriched microbial mRNA was prepared for sequencing using TruSeq Stranded mRNA Library Prep kit (Illumina). Subsequently, the library was sequenced using an Illumina HiSeq 2500 (101 bp paired end sequencing). 118 million paired-end reads were generated and are deposited under the ENA study `PRJNA419191`.

As part of the previous work, reads were partitioned with `khmer`, assembled with `Velvet` and proteins were predicted and annotated with Enzyme Commission (EC) numbers using `MGKit` with the `Uniprot` database.

### Recovery of haplotypes with Gretel

To recover industrially relevant enzyme isoforms from the metatranscriptome, we focused our attention on hydrolases known to be found in the rumen [1]. The existing GFF was filtered to create a subset of all entries with Enzyme Commission (EC) numbers 3.2, 3.4 and 5.3. 3,419 regions from the GFF were identified and were cross-referenced to the new read alignment. Regions were filtered with the criteria:

- minimum coverage *≥* mean minimum coverage (19.7*×*)
- length *≥* new mean region length (615.7 nt)
- standard deviation of coverage *≤* average standard deviation of coverage over remaining regions (76.79*×*)

Filtering returned 259 possible candidates. Each sample’s original short-reads were re-aligned to the existing assembly with `bowtie2` (`--local`) before merging all samples with `samtools merge` to create one canonical alignment of all reads (248,092,426 alignments). `Gretel` was individually executed over the 259 regions, using the aligned reads to recover haplotypes.

Haplotyping was possible at 257 loci. To empirically verify our haplotype predictions, forward and reverse primers were generated with `pd5` [44] using the DNA sequences of the recovered haplotype as a template.

Each set of recovered haplotypes was sorted by descending likelihood. For each haplotype, a corresponding “flattened” consensus was calculated by flipping any base that disagreed with the base call of any haplotype with a better likelihood, to an ‘N’. `pd5` [44] was executed on each consensus with the goal to find a forward and reverse primer that covered the most number of recovered haplotypes, whilst attempting to keep the selected template region as long as possible. Primers could be between 25 and 40 nt, with an annealing temperature between 55 and 65°C. For laboratory analysis, 10 regions were hand-selected (and *ThermoFisher Custom Value Oligos* were synthesized) considering the criteria:

- gene length
- primer template length
- number of predicted haplotypes
- distribution of haplotype likelihoods
- evidence of similar gene sequence in databases
- number of haplotypes that could be captured by generated primers

### PCR Amplicons

Stock RNA from the 32 samples was pooled in equal concentration. Gene-specific reverse transcription for the ten chosen genes (Table 2) was performed with a *Qiagen QuantiTectQ*® *Kit*. Thus, each selected region had an individual corresponding cDNA library.

Gene-specific PCR (30 cycles, 65°C annealing temperature for 20s) was performed for each of the 10 genes with *New England Biolabs PhusionQ*® *High-Fidelity DNA Polymerase*, using the corresponding cDNA (1:10 dilution) and primer pair. Bands were excised following gel electrophoresis and DNA extracted with a *Qiagen QIAquickQ*® *Gel Extraction Kit*. PCR, gel electrophoresis and extraction were repeated to manufacture a sufficient number of amplicons for PacBio sequencing.

Four of the 10 sequences (G11, G31, G90 and G123) could be produced at the expected length and adequate amount for sequencing. Isolated DNA was verified via Sanger sequencing at the Translational Genomics Facility, Aberystwyth. G11 was contaminated with rRNA carryover from the reverse transcription and no haplotypes could be determined.

### PacBio Sequencing

The remaining three amplicons (G31, G123 and G90) were sequenced via PacBio circular consensus sequencing (CCS) on the Sequel platform at the Centre for Genomic Research at Liverpool. Sequencing generated 10,510,553 subreads. CCS reads were constructed with pbccs (`SMRTLink v6.0`) with parameters `--minReadScore=0.65 --minLength=100 --minPasses=20`, yielding 149,003 consensus sequences. The CCS BAM is available from ENA via accession `ERR3446841`. The CCS reads were extracted as FASTQ with bamtools convert and aligned against the set of `Gretel` haplotypes with `minimap2` (`-ax map-pb --MD --secondary=no`), thereby matching each CCS read to its closest `Gretel` haplotype.

### Haplotype Verification

To determine how well the CCS reads supported the `Gretel` haplotypes, the alignments were parsed with our `haplify_zmw.py` script, available from our `gretel-test` repository. Each position along an aligned CCS read was compared to the `Gretel` haplotype, if that position was previously determined a SNP by `gretel-snpper` when the haplotypes were initially recovered. For each CCS read, the script counts the number of matches and mismatches (that is, the number of SNP positions on the CCS read that do and do not match the `Gretel` haplotype, respectively). Bases on a CCS read with a quality score below Q30 were masked to remove noise. Although circular consensus sequencing is expected to give a very high level of accuracy (we observe an average per-base Q-score of 92.1), the CCS reads contained indels when aligned to the `Gretel` haplotypes. These indels are unexpected as they do not appear in the corresponding Illumina data. For this reason, the parsing script also ignored any insertions with respect to the `Gretel` haplotype.

### Hansel and Gretel reveals the landscape of a pangenome

As part of a previous study to explore ruminant methanogenesis, conducted by Shi *et al.* [45], the rumen of 10 rams was characterised through deep whole-genome metagenomic shotgun sequencing. Samples were sequenced with an Illumina HiSeq 2000 (2 *×* 150 bp), generating approximately 50 GB of reads per sample (1020 GB). Metagenomic reads for two samples of the sheep microbiome were downloaded from the European Nucleotide Archive (`SRR873595` and `SRR873610`). 14 Hungate [46] reference sequences for *R. flavefaciens* were selected.

The Hungate references were annotated as part of a now published work using `CowPI` [47], yielding a GFF of annotations for each of the reference genomes. 16,801 CDS regions were annotated across the 14 reference genomes (Supplementary Table 4).

Reads from both samples were aligned to each of the 14 chosen Hungate references separately with `bowtie2` (`--sensitive-local`), producing 28 BAM files. The intuition behind allowing a single read to map to multiple references was to overcome the potential for some regions of the union of references to act as a sink (and soak up the majority of the reads). This allows a read to contribute evidence to haplotypes on more than one reference, but arguably is a fairer strategy than allowing a read to only contribute to a single region of a single genome. A total of 29,476,758 short reads aligned to the references.

Variant calling was conducted as before with `gretel-snpper`, generating a VCF file for every contig in each of the two samples alignment BAMs. Haplotyping was then carried out in parallel with `Gretel` on every combination of sample BAM and reference GFF annotation (n=33602).

For each GFF annotation, we pooled the corresponding haplotypes from the two different sample runs to create 16,801 bags of haplotypes. As `Gretel` does not yet provide automated decision making for quality control of haplotypes, we employed conservative quality control; where there were 4 or more haplotypes returned for a region, haplotypes with likelihoods lower than *−*1000 (the same arbitrary cutoff used to filter out the very worst HIV haplotypes in Supplementary Section 13) were discarded. From the remaining haplotypes the top quartile were selected for downstream analysis. `Gretel` produced 119,151 haplotypes for downstream analysis.

`eggnog-mapper` was used to confirm the reading frame and assign an EggNOG gene family (“NOG”) for each of the haplotype bags. To assess differences in variation observed between the haplotypes from different gene families in a manner that is robust to read coverage and underlying population size, we used the dN/dS ratio, typically used to quantify adaptive evolution between species [48].

For all regions with more than two haplotypes, pairwise dN/dS comparisons were performed by `CRANN` [49] (providing each region’s haplotype set as a FASTA to `CRANN`’s interactive shell via Menu Option 1, and calculating pairwise distances with default parameters via Menu Option 5). Ratios from all pairs were averaged for each of the 16,801 regions. Average dN/dS ratios were then grouped by their EggNOG gene familiy identifier (n=2135), and the average was calculated, allowing for comparison of dN/dS ratios between haplotypes assigned to different EggNOG gene families. Additionally to infer the core and accessory genome, for each EggNOG gene family, we counted the number of references (out of 14) that featured at least one `emapper` annotation of that family.

## Ethics approval and consent to participate

No ethical approval was necessary for this study.

## Availability of data and materials

Our `Hansel` and `Gretel` framework is freely available, open source software available online at https://github.com/samstudio8/Hansel and https://github.com/samstudio8/gretel, respectively.

The code used to generate metahaplomes and synthetic reads for both the randomly generated and real-gene haplotypes, and the testing data used to evaluate our methods is also available online via https://github.com/samstudio8/gretel-test. PacBio sequence data used to haplotype verification are available via ENA study `PRJEB23483`. RNA metatranscriptome data are available via ENA study `PRJNA419191`.

## Author’s contributions

SN, AC, CJC, WA with collaboration from KG and LS discussed and defined the theoretical problem. SN, CJC, AC and WA chose data and designed *in silico* experiments. SN wrote the code and documentation and executed experiments. *In vitro* experiments were designed by SN and WA with collaboration from AE and AS. SH provided rumen metatranscriptome RNA and Illumina sequencing. SN performed laboratory work under the supervision of WA. SN analyzed and interpreted the results with AC, CJC and WA. All authors contributed to the manuscript.

## Acknowledgements

SN wishes to acknowledge Francesco Rubino for his prior work on the assembly and annotation of the rumen metatranscriptome data, Toby Wilkinson for allowing us to use his work on annotation of the Hungate collection prior to its publication, Martin Pollard (Wellcome Sanger Institute) for useful discussion on the PacBio sequencing technology and Nicholas Loman (University of Birmingham) for comments and suggestions on the manuscript. SN and WA acknowledge John Kenny and Margaret Hughes at the Centre for Genomic Research at Liverpool.

## Additional Files

**Additional file 1 — Supplementary Materials**

Mathematical definition of the **metahaplome**, the `Hansel` graph and equations used by `Gretel` and additional results.

## References

[1] Rubino, F., Carberry, C., Waters, S.M., Kenny, D., McCabe, M.S., Creevey, C.J.: Divergent functional isoforms drive niche specialisation for nutrient acquisition and use in rumen microbiome. The ISME Journal (2017)

[2] Callahan, B.J., McMurdie, P.J., Holmes, S.P.: Exact sequence variants should replace operational taxonomic units in marker-gene data analysis. The ISME Journal, 2017119 (2017)

[3] Zhang, C., Kim, S.-K.: Research and application of marine microbial enzymes: status and prospects. Marine Drugs 8(6), 1920–34 (2010). doi:10.3390/md8061920

[4] Kuleshov, V., Jiang, C., Zhou, W., Jahanbani, F., Batzoglou, S., Snyder, M.: Synthetic long-read sequencing reveals intraspecies diversity in the human microbiome. Nature Biotechnology 34, 64–69 (2016). doi:10.1038/nbt.3416

[5] Pulido-Tamayo, S., Sánchez-Rodríguez, A., Toon Swings, T., Van den Bergh, B., Dubey, A., Steenackers, H., Michiels, J., Fostier, J., Marchal, K.: Frequency-based haplotype reconstruction from deep sequencing data of bacterial populations. Nucleic Acids Res 43(16), 105 (2015). doi:10.1093/nar/gkv478

[6] Baaijens, J.A., El Aabidine, A.Z., Rivals, E., Schönhuth, A.: De novo assembly of viral quasispecies using overlap graphs. Genome Research 27, 835–848 (2017)

[7] Quince, C., Delmont, T.O., Raguideau, S., Alneberg, J., Darling, A.E., Collins, G., Eren, A.M.: Desman: a new tool for de novo extraction of strains from metagenomes. Genome Biology 18(1), 181 (2017). doi:10.1186/s13059-017-1309-9

[8] Luo, C., Knight, R., Siljander, H., Knip, M., Xavier, D. R. J. & Gevers: ConStrains identifies microbial strains in metagenomic datasets. Nature Biotechnology 33, 1045–105 (2015)

[9] Handelsman, J., Rondon, M.R., Brady, S.F., Clardy, J., Goodman, R.M.: Molecular biological access to the chemistry of unknown soil microbes: a new frontier for natural products. Chemistry & biology 5(10), 245–249 (1998)

[10] Cilibrasi, R., Van Iersel, L., Kelk, S., Tromp, J.: On the complexity of several haplotyping problems. In: Algorithms in Bioinformatics, pp. 128–139. Springer, ??? (2005)

[11] Lancia, G., Bafna, V., Istrail, S., Lippert, R., Schwartz, R.: SNPs problems, complexity, and algorithms. In: Algorithms—ESA 2001, pp. 182–193. Springer, ??? (2001)

[12] Edge, P., Bafna, V., Bansal, V.: HapCUT2: robust and accurate haplotype assembly for diverse sequencing technologies. Genome Research (2016)

[13] Aguiar, D., Istrail, S.: HapCompass: a fast cycle basis algorithm for accurate haplotype assembly of sequence data. Journal of Computational Biology 19(6), 577–590 (2012)

[14] Motazedi, E., Finkers, R., Maliepaard, C., de Ridder, D.: Exploiting next-generation sequencing to solve the haplotyping puzzle in polyploids: a simulation study. Briefings in Bioinformatics, 126 (2017)

[15] Aguiar, D., Istrail, S.: Haplotype assembly in polyploid genomes and identical by descent shared tracts. Bioinformatics 29(13), 352–360 (2013)

[16] Lancia, G.: Algorithmic approaches for the single individual haplotyping problem. RAIRO-Operations Research 50(2), 331–340 (2016)

[17] Namiki, T., Hachiya, T., Tanaka, H., Sakakibara, Y.: MetaVelvet: An extension of Velvet assembler to de novo metagenome assembly from short sequence reads. Nucleic Acids Res 40(20), 155 (2012)

[18] Boisvert, S., Raymond, F., Godzaridis, É., Laviolette, F., Corbeil, J.: Ray Meta: scalable de novo metagenome assembly and profiling. Genome Biology 13(12), 122 (2012)

[19] Vollmers, J., Wiegand, S., Kaster, A.-K.: Comparing and evaluating metagenome assembly tools from a microbiologist’s perspective-Not only size matters! PLoS One 12(1), 0169662 (2017)

[20] Sieber, C.M., Probst, A.J., Sharrar, A., Thomas, B.C., Hess, M., Tringe, S.G., Banfield, J.F.: Recovery of genomes from metagenomes via a dereplication, aggregation and scoring strategy. Nature microbiology, 1 (2018)

[21] Quince, C., Walker, A.W., Simpson, J.T., Loman, N.J., Segata, N.: Shotgun metagenomics, from sampling to analysis. Nature Biotechnology 35(9), 833–844 (2017)

[22] Schloissnig, S., Arumugam, M., Sunagawa, S., Mitreva, M., Tap, J., Zhu, A., Waller, A., Mende, D.R., Kultima, J.R., Martin, J., et al.: Genomic variation landscape of the human gut microbiome. Nature 493(7430), 45–50 (2013)

[23] Consortium, H.M.P., et al.: Structure, function and diversity of the healthy human microbiome. Nature 486(7402), 207–214 (2012)

[24] Baaijens, J.A., Van der Roest, B., Köster, J., Stougie, L., Schönhuth, A.: Full-length de novo viral quasispecies assembly through variation graph construction. Bioinformatics (2019). doi:10.1093/bioinformatics/btz443

[25] Baaijens, J.A., Stougie, L., Schönhuth, A.: Viral quasispecies reconstruction via contig abundance estimation in variation graphs. bioRxiv (2019). doi:10.1101/645721

[26] Prabhakaran, S., Rey, M., Zagordi, O., Beerenwinkel, N., Roth, V.: HIV haplotype inference using a propagating Dirichlet process mixture model. IEEE/ACM Trans. Comput. Biol. Bioinform., 182–191 (2013)

[27] Zagordi, O., Bhattacharya, A., Eriksson, N., Beerenwinkel, N.: ShoRAH: estimating the genetic diversity of a mixed sample from next-generation sequencing data. BMC bioinformatics 12(1), 119 (2011)

[28] Ahn, S., Vikalo, H.: Joint haplotype assembly and genotype calling via sequential Monte Carlo algorithm. BMC Bioinformatics 16(1), 223 (2015)

[29] Töpfer, A., Zagordi, O., Prabhakaran, S., Roth, V., Halperin, E., Beerenwinkel, N.: Probabilistic inference of viral quasispecies subject to recombination. Journal of Computational Biology 20(2), 113–123 (2013)

[30] Kuleshov, V.: Probabilistic single-individual haplotyping. Bioinformatics 30(17), 379–85 (2014). doi:10.1093/bioinformatics/btu484

[31] Jayasundara, D., Saeed, I., Maheswararajah, S., Chang, B.C., Tang, S.-L., Halgamuge, S.K.: ViQuaS: an improved reconstruction pipeline for viral quasispecies spectra generated by next-generation sequencing. Bioinformatics 31(6), 886–96 (2015)

[32] Dagan, T., Martin, W.: The tree of one percent. Genome biology 7(10), 118 (2006)

[33] Segata, N., Waldron, L., Ballarini, A., Narasimhan, V., Jousson, O., Huttenhower, C.: Metagenomic microbial community profiling using unique clade-specific marker genes. Nature Methods 8, 811–814 (2012)

[34] Wood, D.E., Salzberg, S.L.: Kraken: ultrafast metagenomic sequence classification using exact alignments. Genome biology 15(3), 46 (2014)

[35] Dilthey, A., Jain, C., Koren, S., Phillippy, A.M.: Metamaps – strain-level metagenomic assignment and compositional estimation for long reads. bioRxiv (2018). doi:10.1101/372474

[36] Kolmogorov, M., Yuan, J., Lin, Y., Pevzner, P.A.: Assembly of long, error-prone reads using repeat graphs. Nature biotechnology 37(5), 540 (2019)

[37] Langmead, B., Salzberg, S.: Fast gapped-read alignment with Bowtie 2. Nature Methods 9, 357–359 (2012)

[38] Li, H., Handsaker, B., Wysoker, A., Fennell, T., Ruan, J., Homer, N., Marth, G., Abecasis, G., Durbin, R., 1000 Genome Project Data Processing Subgroup: The sequence alignment/map (SAM) format and SAMtools. Bioinformatics 25, 2078–9 (2009)

[39] DePristo, M., Banks, E., Poplin, R., Garimella, K., Maguire, J., Hartl, C., Philippakis, A., del Angel, G., Rivas, M.A., Hanna, M., McKenna, A., Fennell, T., Kernytsky, A., Sivachenko, A., Cibulskis, K., Gabriel, S., Altshuler, D., Daly, M.: A framework for variation discovery and genotyping using next-generation DNA sequencing data. Nature Genetics 43, 491–498 (2011)

[40] Rambaut, A., Grass, N.C.: Seq-Gen: an application for the Monte Carlo simulation of DNA sequence evolution along phylogenetic trees. Computer applications in the biosciences: CABIOS 13(3), 235–238 (1997). doi:10.1093/bioinformatics/13.3.235

[41] Edgar, R.C.: MUSCLE: a multiple sequence alignment method with reduced time and space complexity. BMC Bioinformatics 5(1), 113 (2004)

[42] Li, D., Liu, C.-M., Luo, R., Sadakane, K., Lam, T.-W.: MEGAHIT: an ultra-fast single-node solution for large and complex metagenomics assembly via succinct de Bruijn graph. Bioinformatics 31(10), 1674–1676 (2015). doi:10.1093/bioinformatics/btv033

[43] Huws, S.A., Edwards, J.E., Creevey, C.J., Stevens, P.R., Lin, W., Girdwood, S.E., Pachebat, J.A., Kingston-Smith, A.H.: Temporal dynamics of the metabolically active rumen bacteria colonizing fresh perennial ryegrass. FEMS Microbiology Ecology 92(1), 137 (2016)

[44] Riley, M.C., Aubrey, W., Young, M., Clare, A.: Pd5: A general purpose library for primer design software. PloS one 8(11), 80156 (2013)

[45] Shi, W., Moon, C.D., Leahy, S.C., Kang, D., Froula, J., Kittelmann, S., Fan, C., Deutsch, S., Gagic, D., Seedorf, H., et al.: Methane yield phenotypes linked to differential gene expression in the sheep rumen microbiome. Genome research 24(9), 1517–1525 (2014)

[46] Seshadri, R., Leahy, S.C., Attwood, G.T., Teh, K.H., Lambie, S.C., Cookson, A.L., Eloe-Fadrosh, E.A., Pavlopoulos, G.A., Hadjithomas, M., Varghese, N.J., et al.: Cultivation and sequencing of rumen microbiome members from the hungate1000 collection. Nature biotechnology 36(4), 359(2018)

[47] Wilkinson, T.J., Huws, S.A., Edwards, J.E., Kingston-Smith, A., Siu Ting, K., Hughes, M., Rubino, F., Friedersdorff, M., Creevey, C.: Cowpi: a rumen microbiome focussed version of the picrust functional inference software. Frontiers in microbiology 9, 1095 (2018)

[48] Li, W.-H.: Unbiased estimation of the rates of synonymous and nonsynonymous substitution. Journal of molecular evolution 36(1), 96–99 (1993)

[49] Creevey, C., McInerney, J.O.: Crann: detecting adaptive evolution in protein-coding dna sequences. Bioinformatics 19(13), 1726–1726 (2003)

