## Supplementary Materials for "Recovery of gene haplotypes from a metagenome"

#### Contents

**1 The metahaplome** **2**

**2 Hansel as a graph** **3**

**3 Probabilistic edge weights** **3**

**4 Simplification of conditional edge weights** **4**

**5 Estimation of probabilities** **4**

**6 Smoothing** **5**

**7 Reweighting** **5**

**8 Stopping criterion** **5**

**9 Haplotype scoring** **5**

**10 Hansel and Gretel is capable of recovering haplotypes from mixed strains** **6**

**11 Hansel and Gretel recovers gene haplotypes from a real metagenome** **7**

### 1 The metahaplome

We formulate the concept of the metahaplome, and the problem of recovering haplotypes from a microbial community formally. First we must provide the following definitions:

- $\Omega$
  An environment of microbial organisms.
- $O$
  A set[1] containing each full genomic sequence, of each individual organism in environment $\Omega$.
  $O$ is *the* metagenome of $\Omega$: encompassing all possible genomes in the environment.
- $o[i:j]$
  A sub-sequence $i..j$ of some genome $o \in O$.
- Gene $g$
  A known DNA sequence $g$, potentially responsible for the production of a protein capable of performing a catalytic reaction of interest, such as the hydrolysis of cellulose
- $\Delta(s, g)$
  Any function $\Delta$ that can determine whether a DNA sequence $s$ (such as a sufficiently sized sub-sequence $o[i:j]$) has sufficient sequence similarity to $g$, such as `BLAST`. $\Delta$ may return a boolean, or a real value (*e.g.* expect value) that can be coerced to a boolean via a user-selected threshold.
- $\Gamma_g = set(\wr o_k[i:j] \mid \Delta(o_k[i:j], g),\ k \in 1..|O|,\ i, j \in 1..|o_k|,\ i < j, j - i \approx |g| \int)$
  One may collect a bag of sub-sequences across the elements of $O$, determined to have sequence similarity to $g$ by $\Delta$. $\Gamma_g$ represents the set of unique sequences contained in the bag.

We define $\Gamma_g$ as the **metahaplome** for the gene $g$, in the metagenome $O$. Consider $g$ encodes a protein that performs some biological function of interest, then each $\gamma \in \Gamma_g$ is a **gene haplotype** of $g$. That is, generally, $\gamma$ is a DNA sequence that encodes an isoform of $g$. Ideally, we wish to recover the set of all such haplotypes: $\Gamma_g$. Of course, the metagenome $O$ is unknown, and our insight to $O$ is obtained through environmental sampling and DNA sequencing. We must recover haplotypes from the available evidence, and require the following additional definitions to define the recovery of gene haplotypes from a metagenome:

- $\sigma = \text{Sample}(\Omega)$
  - A sample taken from microbial environment $\Omega$.
  - $\sigma \subset \Omega$
- $M$
  - A set[2] containing the genomes $m \in M$, for each individual captured in the sample $\sigma$.
  - $M$ represents the metagenome that was captured in the sample $\sigma$, and is our insight into $O$.
  - $M$ is not necessarily or likely to be representative of the entire metagenome $O$.
  - $M \subset O$
- $R = \text{Seq}(\sigma)$
  - $R$ is the set of **reads** obtained from the sequencing of isolated DNA from sample $\sigma$.
  - A read consists of a sequence of nucleotide bases $r_i[j] \in \{A, C, G, T, N\}, i \in 1..|R|, j \in 1..|r_i|$.
  - A read describes a fragment of some genome $m_k[u:v] \in M$, with some degree of error.
  - Due to sampling bias acquiring $\sigma$ from $\Omega$, $R$ is unlikely to be representative of the true genetic diversity in $O$.
  - Additionally, due to sequencing and PCR biases and error, $R$ is unlikely to provide uniform and non-zero coverage of the residues across all $m \in M$.
- $C = \text{Assemble}(R)$
  - Contig set $C$ (**Assembly**) constructed *de novo* from the reads $R$ by some `Assemble` operation.
  - $c_i[j] \in \{A, C, G, T, N\}$ for $i \in 1..|C|, j \in 1..|c_i|$.
  - `Assemble` attempts to reconstruct $M$ from the reads $R$, but typically fails to distinguish between highly similar sequences that should create distinct $c \in C$.
  - $C$ poses as a **pseudo-reference** for the metagenome $M$.
  - Alternatively, $C$ could be a set of high-quality reference sequences, if available.

[1] Arguably one could also consider this definition as a bag, where we could have duplicates of genomes in the environment. However, the later set builder definitions are less cluttered by restricting this to a unique set.

[2] Again, we refer to this as a set, rather than a bag.

- $A = \text{Align}(R, C)$
  - **Alignment** $A$, generated by aligning read set $R$ to contig set $C$ with operation `Align`.
  - $A_{c_k[i:j]}$ is the set of read alignments in $A$ that cover any position between $i$ and $j$ on $c_k \in C$.
- $S = \text{Call}(A_{c_k})$
  - The set of genomic positions on a contig $c_k \in C$ determined to be single nucleotide polymorphisms (SNPs) by the operation `Call`, given $A_c$: the alignments of $R$ against $c_k$.
  - `Call` may simply consider each 'column' $A_{c_k}[i]$ for $i \in 1..|c_k|$ and determine position $i$ as a variant if there is a disagreement on the nucleotide at that position across the aligned reads. `Call` may also be a more complex variant prediction algorithm.

Our goal is to determine the metahaplome $\Gamma_g$, for some gene of interest $g$. Although $M$ will not be entirely representative of $O$, it is the only evidence of the sequence diversity available to us. Thus we adjust our goal to instead find the most likely elements of $\Gamma_g$, given the evidence that can be derived from $M$, via the alignments $A$ and SNP sites $S$.

#### 2 Hansel as a graph

Consider an alphabet of symbols, $\Sigma$ (*e.g.* $\{A, C, G, T, N, -\}$) and a list of $n$ SNP positions $1..n$. Symbols $\varnothing_S$ and $\varnothing_E$ represent special sentinal positions at the start and end of the SNP positions (0 and $n+1$ respectively). As described in our article, the `Hansel` structure $H$ can be considered as a graph $G = (V, E)$. Here, we define $V$, and $E$:

$$E = \bigcup_{i=1..n} \{(A_i, B_{i+1}) \mid H[A, B, i, i+1] > 0, A \in (\Sigma \cup \varnothing_S), B \in (\Sigma \cup \varnothing_E)\} \tag{S1}$$

$$V = \{v \mid (v, w) \in E\} \cup \{v \mid (w, v) \in E\} \tag{S2}$$

$E$ represents the set of edges, where an edge $(A_i, B_{i+1})$ is determined to exist in $E$ if there exists at least one read whereby symbol $A$ was observed at position $i$ to co-occur with symbol $B$ at SNP position $i+1$.

It should be noted, that although $G$ can be constructed from $H$ such that it is undirected and contains cycles, both properties lead to nonsensical haplotypes. Under such circumstances, `Gretel` could construct a path that visits multiple nodes that appear at the same $i$, or a trail that visits the same node multiple times. Such sequences would be meaningless in the context of haplotype construction, thus the interface to `Hansel` acts in such a way that $G$ is a directed, acyclic graph.

We can define a haplotype as an alternating sequence of nodes ($v \in V$) and edges ($e \in E$). A path must always start and end at the special sentinel symbols $\varnothing_S$ and $\varnothing_E$, respectively.

$$\hat{h} = \varnothing_S, e_0, v_1, e_1, v_2, e_2, ..., v_{n-1}, e_n, v_n, e_{n+1}, \varnothing_E \tag{S3}$$

Although, as only one directed edge between some $v_i$ and $v_{i+1}$ may exist, we can define $\hat{h}$ as a sequence of $v \in V$:

$$\hat{h} = \varnothing_S, v_1, v_2, ..., v_{n-1}, v_n, \varnothing_E \tag{S4}$$

#### 3 Probabilistic edge weights

However, if the construction of $G$ does not consider elements in $H[A, B, i, j]$ where $abs(i-j) > 1$ (non-adjacent SNPs) it is likely one will recover haplotypes that do not actually exist.

Given the pairwise information available in $H$, for both adjacent, and non-adjacent SNPs, across all reads, edges in the graph $G$ derived from $H$ can be weighted probabilistically. We attempt to determine the next most likely symbol in a sequence, considering both the marginal distribution of symbols at the next position and the likelihood of those symbols appearing next, given an already observed partial sequence. That is, the next symbol $v_{i+1}$ in a path depends not only on the current symbol ($v_i$) but some number of previous symbols ($v_{i-1}, v_{i-2}...v_0$).

The outgoing edges from $v_i$ are probabilistically weighted by exploiting the observations stored in the `Hansel` structure to create probabilities. These probabilities then determine the likelihood of moving from some $v_i$ to each of the possible $v_{i+1}$.

We take a Bayesian approach to the problem of probabilistically weighting edges in `Hansel`'s graph representation. We define the probability of selecting $v_{i+1}$, conditioned on the path observed so far:

$$\begin{aligned}
&\mathbb{P}(v_{i+1} \mid v_1, v_2, ..., v_{i-1}, v_i) \\
&\quad \propto \mathbb{P}(v_1, v_2, ..., v_i, v_{i+1}) \\
&\quad = \mathbb{P}(v_1 \mid v_2...v_{i+1}) \times \mathbb{P}(v_2, ...v_{i+1}) \\
&\quad = \mathbb{P}(v_1 \mid v_2...v_{i+1}) \times \mathbb{P}(v_2 \mid v_3...v_{i+1}) \times \mathbb{P}(v_3, ...v_{i+1}) \\
&\quad = \mathbb{P}(v_1 \mid v_2...v_{i+1}) \times \mathbb{P}(v_2 \mid v_3...v_{i+1}) \times ... \times \mathbb{P}(v_{i-1} \mid v_i, v_{i+1}) \\
&\quad\quad \times \mathbb{P}(v_i \mid v_{i+1}) \times \mathbb{P}(v_{i+1})
\end{aligned} \tag{S5}$$

### 4 Simplification of conditional edge weights

Clearly, the number of factors in Equation S5 increases with $i$. For longer paths (more single nucleotide polymorphisms detected along the target region of interest), evaluating the equation becomes more computationally expensive, and risks potentially compounding estimation errors.

To construct a whole path $p$ from $v_1...v_n$, the upper bound for the number of iterations will be $|\Sigma| \times n$ with calculations becoming increasingly complex as $i$ increases.

To reduce complexity, we make an assumption of conditional independence between variants. Whilst this seems counter intuitive, the Naive Bayes model can deliver robust results despite its coarse assumption.

Thus we may simplify our previous equation and consider only the pairwise appearances of each $v_i$ encountered thus far against $v_{i+1}$.

$$\begin{aligned}
&\mathbb{P}(v_{i+1} \mid v_1, v_2, ..., v_{i-1}, v_i) \\
&\quad \approx \mathbb{P}(v_{i+1}) \times \mathbb{P}(v_1 \mid v_{i+1}) \times \mathbb{P}(v_2 \mid v_{i+1}) \times ... \\
&\quad = \mathbb{P}(v_{i+1}) \prod_{j=1}^{i} \mathbb{P}(v_j \mid v_{i+1})
\end{aligned} \tag{S6}$$

However as discussed in our article, individual reads will not cover all SNP positions $1..n$ (if they did, we would not have to define this problem). Thus, we need not consider all variants in the current path when evaluating edge weights. Instead, we could limit the number of variants to consider, from the current position in the path $i$, back some small and sensible number of steps $L$:

$$\mathbb{P}(v_{i+1} \mid v_{i-L}, ..., v_{i-2}, v_{i-1}, v_i) = \mathbb{P}(v_{i+1}) \prod_{l=0}^{L-1} \mathbb{P}(v_{i-l} \mid v_{i+1}) \tag{S7}$$

Additionally, to overcome inaccuracies encountered through floating point error when performing mathematical operations on very small decimals, `Gretel` uses log probabilities instead. Via the log identity $log(ab) = log(a) + log(b)$ the product of the conditional probabilities becomes a sum of the log conditional probabilities:

$$log_{10}(\mathbb{P}(v_{i+1} \mid v_{i-L}, ..., v_{i-2}, v_{i-1}, v_i)) = log_{10}(\mathbb{P}(v_{i+1})) + \sum_{l=0}^{L-1} log_{10}(\mathbb{P}(v_{i-l} \mid v_{i+1})) \tag{S8}$$

We define $L$ as the the 'lookback' size, the number of variants of the current path to consider when selecting $v_{i+1}$. Conveniently, there is a reasonable intuition available for selecting a value for $L$: the mean number of SNP sites covered by the observed reads. Thus we avoid the scenario of introducing an algorithmically influential but difficult to optimize parameter, such as $k$-mer size for metagenomic assembly.

### 5 Estimation of probabilities

Equation S9 provides an estimate for the marginal distribution of a symbol $\beta$ appearing at position $j$.

$$\begin{aligned}
&\hat{\mathbb{P}}(v_j = \beta) \\
&\quad = \frac{\text{Number of reads with symbol } \beta \text{ at position } j}{\text{Number of reads spanning position } j} \\
&\quad = \frac{\sum_{\gamma \in \Sigma} H[\beta,\ \gamma,\ j,\ j+1]}{\sum_{\gamma \in \Sigma} \sum_{\delta \in \Sigma} H[\gamma,\ \delta,\ j,\ j+1]}
\end{aligned} \tag{S9}$$

Equation S8 provides an estimate for the conditional distribution of symbol $\alpha$ appearing at position $i$ given that $\beta$ was observed at position $j$.

$$\begin{aligned}
&\hat{\mathbb{P}}(v_i = \alpha \mid v_j = \beta) \\
&\quad = \frac{\text{Number of reads featuring } \alpha \text{ at } i \text{ and } \beta \text{ at } j}{\text{Number of reads spanning } i \text{ featuring symbol } \beta \text{ at } j} \\
&\quad = \frac{H[\alpha,\ \beta,\ i,\ j]}{\sum_{\gamma \in \Sigma} H[\gamma,\ \beta,\ i,\ j]}
\end{aligned} \tag{S10}$$

#### 6 Smoothing

To avoid the potential of dividing by 0 when using Equation S10 in cases where a suitable read spanning $i$ and $v_j = \beta$ does not exist, we apply Laplace smoothing to effectively add a dummy support read. Future work will investigate alternative smoothing methodology.

$$\begin{aligned}
&\hat{\mathbb{P}}(v_i = \alpha \mid v_j = \beta) \\
&= \frac{1 + \text{Number of reads featuring } \alpha \text{ at } i \text{ and } \beta \text{ at } j}{\text{Variants at } i + \text{Number of reads spanning } i \text{ featuring symbol } \beta \text{ at } j} \\
&= \frac{1 \ + \ H[\alpha,\ \beta,\ i,\ j]}{|\{\gamma_i \mid H[\gamma,\ \sigma, i, i+1] > 0, \gamma \in \Sigma, \sigma \in \Sigma\}| \ + \ \sum_{\gamma \in \Sigma} H[\gamma, \beta, i, j]}
\end{aligned} \tag{S11}$$

#### 7 Reweighting

The paths generated by `Gretel` are probabilistic, but not stochastic. For a given $H$, `Gretel` will always return the same path. After a path has been constructed we can perform some transformation of $H$ to prevent repetitive generation of the same path and return the next most likely path on the next iteration instead.

Given a path $\hat{h}$, we inspect the marginal distribution of each element of the path in order to find the smallest marginal. `Gretel` iterates over each element $\hat{h}[i]$ in the path, and uses the `Hansel` interface to reweight the element $H[\hat{h}[i], \hat{h}[i+1], i, i+1]$ by subtracting the result of multiplying the smallest marginal by the original value for that observation in $H$:

$$\lambda = \min(\{\mathbb{P}(\hat{h}[i]) \mid i = 1..n\}) \tag{S12}$$

$$H[\hat{h}[i],\ \hat{h}[i+1],\ i,\ i+1] = H[\hat{h}[i],\ \hat{h}[i+1],\ i,\ i+1] - (\lambda \times H[\hat{h}[i],\ \hat{h}[i+1],\ i,\ i+1]) \tag{S13}$$

In practice $\lambda$ is capped by `Gretel` in an attempt to stop aggressive reweighting that might otherwise prevent the recovery of closely related haplotypes.

#### 8 Stopping criterion

After multiple iterations of path finding and subsequent reweighting, elements in $H$ will begin to approach 0, causing edges in the graph to become unavailable for traversal. `Gretel` will immediately terminate upon encountering a node in the graph with no viable outgoing edges. That is, the selected symbol at the current $i$ has no non-zero weighted edges to traverse between SNP positions $(i,\ i+1)$ in the graph. Alternatively, if this criterion is not reached after 100 iterations (haplotypes), `Gretel` aborts.

#### 9 Haplotype scoring

`Gretel` can score and rank the haplotypes it recovers. For a completed haplotype, $\hat{h}$, we compute its likelihood based upon the sum of the marginal log probabilities for each element of $\hat{h}$ given the current state of $H$.

$$\hat{h} = \varnothing_S, v_1, v_2, ..., v_{n-1}, v_n, \varnothing_E$$

$$\begin{aligned}
L(\hat{h}) = \mathbb{P}(H \mid \hat{h}) &= \mathbb{P}(v_1 = \hat{h}[1], v_2 = \hat{h}[2], v_3 = \hat{h}[3], ...v_{n-1} = \hat{h}[n-1], v_n = \hat{h}[n]) \\
&= \prod_{i=1}^{n} \hat{\mathbb{P}}(v_i = \hat{h}[i]) \\
&= \prod_{i=1}^{n} \frac{\sum_{\gamma \in \Sigma} H[\hat{h}[i],\ \gamma,\ i,\ i+1]}{\sum_{\gamma \in \Sigma} \sum_{\delta \in \Sigma} H[\gamma,\ \delta,\ i,\ i+1]}
\end{aligned} \tag{S14}$$

To overcome the potential for floating point arithmetic error (Equation S8), we calculate and report the log likelihood.

$$log_{10}(L(\hat{h})) = \sum_{i=1}^{n} log_{10} \left( \frac{\sum_{\gamma \in \Sigma} H[\hat{h}[i],\ \gamma,\ i,\ i+1]}{\sum_{\gamma \in \Sigma} \sum_{\delta \in \Sigma} H[\gamma,\ \delta,\ i,\ i+1]} \right) \tag{S15}$$

### 10 Hansel and Gretel is capable of recovering haplotypes from mixed strains

| | Raw Assembly | $\geq$ 1kbp |
|---|---|---|
| **Contigs** | 17,066 | 6,357 |
| **Total bp** | 67,189,963 | 61,651,258 |
| **Min** | 200 | 1,000 |
| **Average** | 3,937 | 9,698 |
| **Max** | 689,365 | 689,365 |
| **N50** | 53,290 | 63,517 |
| **Time** | 4605 s | - |

Table 1: Statistics for our `MEGAHIT` assembled from read data provided by Quince *et al.*

### 11 Hansel and Gretel recovers gene haplotypes from a real metagenome

#### 11.1 Summary of Gretel's recovered haplotypes

| Amplicon | Haplotype Number | Likelihood | Fewest mismatches (N) | Distinct CCS |
|---|---|---|---|---|
| G123 | 0 | -16.65 | 0 (1592) | 10944 |
| G123 | 2 | -27.29 | 0 (458) | 1859 |
| G123 | 3 | -28.60 | 0 (405) | 7989 |
| G123 | 5 | -32.79 | 0 (351) | 3047 |
| G123 | 8 | -36.02 | 0 (845) | 5306 |
| G123 | 9 | -38.11 | 1 (83) | 1770 |
| G123 | 22 | -45.34 | 1 (167) | 670 |
| G123 | 17 | -69.45 | - | - |
| G123 | 15 | -69.67 | - | - |
| G123 | 16 | -70.20 | - | - |
| G123 | 19 | -70.63 | - | - |
| G123 | 18 | -70.84 | - | - |
| G123 | 14 | -70.86 | - | - |
| G123 | 24 | -73.94 | - | - |
| G123 | 25 | -74.49 | - | - |
| G123 | 28 | -78.97 | - | - |
| G123 | 27 | -79.27 | - | - |
| G123 | 26 | -79.36 | - | - |
| G123 | 30 | -80.07 | - | - |
| G123 | 23 | -80.16 | - | - |
| G123 | 29 | -81.05 | - | - |
| G123 | 32 | -81.24 | - | - |
| G123 | 31 | -82.07 | - | - |
| G123 | 34 | -82.45 | - | - |
| G123 | 66 | -83.89 | - | - |
| G123 | 45 | -83.91 | - | - |
| G123 | 44 | -84.01 | - | - |
| G123 | 46 | -84.74 | - | - |
| G123 | 64 | -84.76 | - | - |
| G123 | 37 | -84.84 | - | - |
| G123 | 67 | -85.34 | - | - |
| G123 | 36 | -85.57 | - | - |
| G123 | 65 | -85.74 | - | - |
| G123 | 71 | -86.04 | - | - |
| G123 | 39 | -86.28 | - | - |
| G123 | 49 | -86.31 | - | - |
| G123 | 43 | -86.35 | - | - |
| G123 | 69 | -86.36 | - | - |
| G123 | 50 | -86.71 | - | - |
| G123 | 70 | -86.99 | - | - |
| G123 | 41 | -87.02 | - | - |
| G123 | 51 | -87.18 | - | - |
| G123 | 73 | -87.18 | - | - |
| G123 | 72 | -87.28 | - | - |
| G123 | 74 | -88.24 | - | - |
| G123 | 75 | -88.49 | - | - |
| G123 | 81 | -89.26 | - | - |
| G123 | 77 | -89.27 | - | - |
| G123 | 80 | -89.55 | - | - |
| G123 | 82 | -89.61 | - | - |
| G123 | 83 | -90.10 | - | - |
| G123 | 60 | -90.16 | - | - |
| G123 | 53 | -90.37 | - | - |
| G123 | 78 | -90.38 | - | - |
| G123 | 59 | -90.61 | - | - |

*Continued on next page...*

...continued from previous page

| Amplicon | Haplotype Number | Likelihood | Fewest mismatches (N) | Distinct CCS |
|---|---|---|---|---|
| G123 | 54 | -90.82 | - | - |
| G123 | 85 | -90.92 | - | - |
| G123 | 62 | -91.47 | - | - |
| G123 | 88 | -91.62 | - | - |
| G123 | 61 | -91.83 | - | - |
| G123 | 56 | -92.39 | - | - |
| G123 | 89 | -92.76 | - | - |
| G123 | 63 | -92.82 | - | - |
| G123 | 91 | -93.42 | - | - |
| G123 | 90 | -93.56 | - | - |
| G123 | 93 | -93.90 | - | - |
| G123 | 92 | -94.45 | - | - |
| G123 | 58 | -95.76 | - | - |
| G123 | 95 | -95.98 | - | - |
| G123 | 97 | -96.80 | - | - |
| G123 | 96 | -96.85 | - | - |
| G123 | 98 | -97.64 | - | - |
| G123 | 99 | -97.86 | - | - |
| G123 | 13 | -159.22 | - | - |
| G123 | 12 | -163.95 | - | - |
| G123 | 11 | -170.75 | - | - |
| G123 | 87 | -179.14 | - | - |
| G123 | 10 | -180.58 | - | - |
| G123 | 7 | -185.15 | - | - |
| G123 | 6 | -228.77 | - | - |
| G123 | 4 | -255.17 | - | - |
| G90 | 0 | -28.13 | 0 (61) | 4543 |
| G90 | 1 | -50.64 | 0 (16) | 2724 |
| G90 | 2 | -66.68 | 3 (6) | 708 |
| G90 | 4 | -75.27 | - | - |
| G90 | 3 | -76.47 | 5 (1) | 309 |
| G90 | 5 | -80.70 | 11 (2) | 1029 |
| G90 | 6 | -88.31 | - | - |
| G90 | 7 | -93.37 | - | - |
| G90 | 13 | -99.84 | 101 (1) | 13 |
| G90 | 10 | -100.93 | 24 (2) | 375 |
| G90 | 11 | -102.27 | 32 (1) | 109 |
| G90 | 8 | -105.21 | 11 (1) | 143 |
| G90 | 9 | -111.07 | 7 (3) | 84 |
| G90 | 12 | -116.73 | 116 (1) | 2 |
| G90 | 14 | -121.15 | - | - |
| G90 | 15 | -125.82 | 46 (1) | 36 |
| G90 | 17 | -126.83 | 35 (1) | 5 |
| G90 | 18 | -130.82 | 130 (1) | 2 |
| G90 | 16 | -133.90 | 21 (1) | 42 |
| G90 | 20 | -134.38 | - | - |
| G90 | 22 | -135.15 | - | - |
| G90 | 25 | -143.28 | - | - |
| G90 | 26 | -145.35 | - | - |
| G90 | 19 | -145.68 | - | - |
| G90 | 23 | -150.75 | - | - |
| G90 | 24 | -152.31 | - | - |
| G90 | 28 | -155.14 | - | - |
| G90 | 27 | -157.49 | 33 (1) | 3 |
| G90 | 30 | -279.10 | - | - |
| G90 | 29 | -317.91 | - | - |
| G90 | 21 | -352.94 | - | - |
| G31 | 0 | -28.29 | 4 (1) | 7804 |
| G31 | 1 | -39.68 | 3 (1) | 8117 |
| G31 | 2 | -46.88 | 8 (3) | 2528 |

Continued on next page...

...continued from previous page

| Amplicon | Haplotype Number | Likelihood | Fewest mismatches (N) | Distinct CCS |
|---|---|---|---|---|
| G31 | 3 | -65.10 | 35 (1) | 4 |
| G31 | 4 | -75.94 | 8 (1) | 1029 |
| G31 | 5 | -82.47 | 19 (1) | 108 |
| G31 | 7 | -96.50 | - | - |
| G31 | 6 | -98.61 | - | - |
| G31 | 11 | -109.13 | 37 (1) | 1 |
| G31 | 10 | -125.10 | - | - |
| G31 | 12 | -152.89 | - | - |
| G31 | 9 | -190.69 | - | - |
| G31 | 8 | -204.00 | - | - |

Table 2: Table enumerating haplotypes recovered by `Gretel` for three amplicons isolated and amplified out of a rumen metatranscriptome. We present each haplotype for the three amplicons, ordered by decreasing likelihood. The last two columns show the fewest number of mismatches to the haploype observed by our `haplify_zmw` script (and how many times that CCS sequence was seen) and the number of unique CCS assigned to that haplotype (including CCS sequences with mismatches). Haplotypes for which there was no PacBio read support are indicated by $-$. `Gretel`'s haplotypes with the best likelihoods for G123 and G90 match those recovered with 0 mismatches. Note the presence of CCS reads with many mismatches to `Gretel` haplotypes (*e.g.* G90#13) indicating potential off-target amplification as part of the PCR process, though these highly mutated CCS sequences are often seen only once (bracketed figures).

##### 11.2 PacBio Circular Consensus Sequence Counts

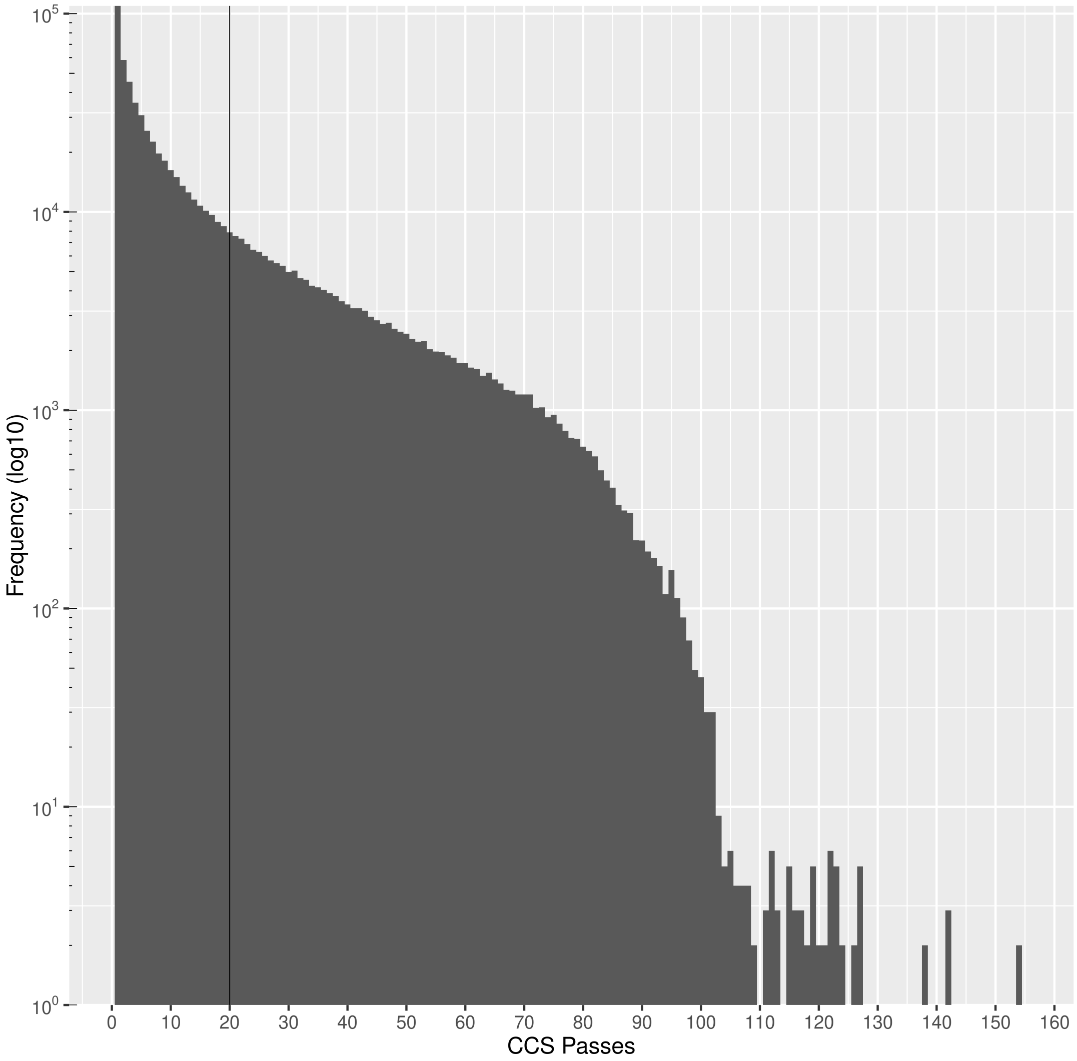

Figure 1: Histogram of log-frequency (y-axis), counting the number of complete sequenced passes (x-axis) for the amplicons sequenced on the Pacific Biosciences Sequel. A vertical line at x=20 delineates the minimum number of passes required for `pbccs` to emit a CCS read for our downstream analysis. 20 passes was chosen to be sure to have enough evidence of the sequence assuming a 15% per-base error rate on the subread.

#### 11.3 CCS reads perfectly match G123 haplotypes

```
BLASTN 2.7.1+

Reference: Zheng Zhang, Scott Schwartz, Lukas Wagner, and Webb
Miller (2000), "A greedy algorithm for aligning DNA sequences", J
Comput Biol 2000; 7(1-2):203-14.

Database: ../../g2019a.full_hap.fasta
           125 sequences; 163,008 total letters

Query= m54118_190409_113909/8848017/ccs
Length=1317

> G123_0__-16.65
Length=1425

 Score = 2433 bits (1317),  Expect = 0.0
 Identities = 1317/1317 (100%), Gaps = 0/1317 (0%)
 Strand=Plus/Minus

Query  1     CGCTTGCAGATGCCTTAGCACCAACGTTGATAGCTACAGGAAGCTCATCTGCTGTCTTCC  60
             ||||||||||||||||||||||||||||||||||||||||||||||||||||||||||||
Sbjct  1402  CGCTTGCAGATGCCTTAGCACCAACGTTGATAGCTACAGGAAGCTCATCTGCTGTCTTCC  1343

Query  61    AGGCTTCATCATCAAACTCACCATCAATAACAGGTGTGCCCTTCTTAACTACTGCAAGTG  120
             ||||||||||||||||||||||||||||||||||||||||||||||||||||||||||||
Sbjct  1342  AGGCTTCATCATCAAACTCACCATCAATAACAGGTGTGCCCTTCTTAACTACTGCAAGTG  1283

Query  121   GCTTAATTACTGTTTCAGCAAAGAACTTGCTGGACTGAGCATGCTTAAATGTTGTGTCGT  180
             ||||||||||||||||||||||||||||||||||||||||||||||||||||||||||||
Sbjct  1282  GCTTAATTACTGTTTCAGCAAAGAACTTGCTGGACTGAGCATGCTTAAATGTTGTGTCGT  1223

Query  181   TAAATACAGCAAGGAAATCACCGTCATTAACTACAATATCAAACTTAACCTTGTTAGCTG  240
             ||||||||||||||||||||||||||||||||||||||||||||||||||||||||||||
Sbjct  1222  TAAATACAGCAAGGAAATCACCGTCATTAACTACAATATCAAACTTAACCTTGTTAGCTG  1163

Query  241   TAAGAGCTTCGAAGTCAACAGGAACATTTACTACTGCCTCGTATCCATTTTCATTTTCTG  300
             ||||||||||||||||||||||||||||||||||||||||||||||||||||||||||||
Sbjct  1162  TAAGAGCTTCGAAGTCAACAGGAACATTTACTACTGCCTCGTATCCATTTTCATTTTCTG  1103

Query  301   TTGCTTCTTCTCTCTTTACTACTACTGATTTGATTCCTGTTCCGTCATCTGCATAAACTG  360
             ||||||||||||||||||||||||||||||||||||||||||||||||||||||||||||
Sbjct  1102  TTGCTTCTTCTCTCTTTACTACTACTGATTTGATTCCTGTTCCGTCATCTGCATAAACTG  1043

Query  361   TAAAGAAATCATCCTGGCCAGTTGTTGTATCTTTTATTACAACTTTAACGTCGATGCCAT  420
             ||||||||||||||||||||||||||||||||||||||||||||||||||||||||||||
Sbjct  1042  TAAAGAAATCATCCTGGCCAGTTGTTGTATCTTTTATTACAACTTTAACGTCGATGCCAT  983

Query  421   TCTCACTCCACATAGGGATAAACTTGCCAGCAACAAACTCATATACATTACCAGCTGAGA  480
             ||||||||||||||||||||||||||||||||||||||||||||||||||||||||||||
Sbjct  982   TCTCACTCCACATAGGGATAAACTTGCCAGCAACAAACTCATATACATTACCAGCTGAGA  923

Query  481   AATCATTATTTACATTCTGTACAAGTGTTATTGACTTTACTTCAGGCTCAAGTTCTCCGG  540
             ||||||||||||||||||||||||||||||||||||||||||||||||||||||||||||
Sbjct  922   AATCATTATTTACATTCTGTACAAGTGTTATTGACTTTACTTCAGGCTCAAGTTCTCCGG  863

Query  541   CATTTGCAATTGCCCAGAAGCAATCCTTAGCCTTGTAGTAATCATCAAAGAGAAGTGGAT  600
             ||||||||||||||||||||||||||||||||||||||||||||||||||||||||||||
Sbjct  862   CATTTGCAATTGCCCAGAAGCAATCCTTAGCCTTGTAGTAATCATCAAAGAGAAGTGGAT  803

Query  601   ACTGTCTTGCGCTTCCGTCTGCGCCGCCACCGTTATTGTTTGATGACTGGAGCCATGAAT  660
             ||||||||||||||||||||||||||||||||||||||||||||||||||||||||||||
Sbjct  802   ACTGTCTTGCGCTTCCGTCTGCGCCGCCACCGTTATTGTTTGATGACTGGAGCCATGAAT  743

Query  661   GCTTATCTGTTACTCCCCAGATTGTCATACCTGTGAAGTTTACGCCGTCTTCTCTAAGTC  720
             ||||||||||||||||||||||||||||||||||||||||||||||||||||||||||||
Sbjct  742   GCTTATCTGTTACTCCCCAGATTGTCATACCTGTGAAGTTTACGCCGTCTTCTCTAAGTC  683

Query  721   TTCTGATTGTGTCATAAACTGCTTTGTATCTCTCCGCAAGTCTGTCATCCTTGGCACTTG  780
             ||||||||||||||||||||||||||||||||||||||||||||||||||||||||||||
Sbjct  682   TTCTGATTGTGTCATAAACTGCTTTGTATCTCTCCGCAAGTCTGTCATCCTTGGCACTTG  623

Query  781   TAGCACCCTTGAAGTCAAGCTCAGTTACCTGAACCTGGTCAACGATAGCTGCATAAGCCT  840
             ||||||||||||||||||||||||||||||||||||||||||||||||||||||||||||
Sbjct  622   TAGCACCCTTGAAGTCAAGCTCAGTTACCTGAACCTGGTCAACGATAGCTGCATAAGCCT  563

Query  841   TGGCTGCTGTCTTGAACTGCTCCATTGAAGGATTGTTGGATGCAATCTGGTAGTGAGCCT  900
             ||||||||||||||||||||||||||||||||||||||||||||||||||||||||||||
Sbjct  562   TGGCTGCTGTCTTGAACTGCTCCATTGAAGGATTGTTGGATGCAATCTGGTAGTGAGCCT  503

Query  901   GCATACCCATACCATCAATTCTTGCACCAGGTGTTGCCTTAACATCTCTAAGAAGCTGGC  960
             ||||||||||||||||||||||||||||||||||||||||||||||||||||||||||||
Sbjct  502   GCATACCCATACCATCAATTCTTGCACCAGGTGTTGCCTTAACATCTCTAAGAAGCTGGC  443

Query  961   AGATACCACCCATCTTGCTGCTTACTGTCTCGTTATAATCGTTATAGAAAAGAGCGATAT  1020
             ||||||||||||||||||||||||||||||||||||||||||||||||||||||||||||
Sbjct  442   AGATACCACCCATCTTGCTGCTTACTGTCTCGTTATAATCGTTATAGAAAAGAGCGATAT  383

Query  1021  CAGCAGGCATATATCTGTTAGCATATACGAATGCATTTGTGATGAACTCCTGTGAGCCGT  1080
             ||||||||||||||||||||||||||||||||||||||||||||||||||||||||||||
Sbjct  382   CAGCAGGCATATATCTGTTAGCATATACGAATGCATTTGTGATGAACTCCTGTGAGCCGT  323

Query  1081  AAACTCCCCACCACTCTGATCTCTCAGATGCATTTCTGTAAGTACCTGTTCCATCACTTA  1140
             ||||||||||||||||||||||||||||||||||||||||||||||||||||||||||||
Sbjct  322   AAACTCCCCACCACTCTGATCTCTCAGATGCATTTCTGTAAGTACCTGTTCCATCACTTA  263

Query  1141  CTGCTTCGTTAACAACATCCCATCCATAGAAGAGATCTTTGTACTTGCTGTCTTCTGCTG  1200
             ||||||||||||||||||||||||||||||||||||||||||||||||||||||||||||
Sbjct  262   CTGCTTCGTTAACAACATCCCATCCATAGAAGAGATCTTTGTACTTGCTGTCTTCTGCTG  203

Query  1201  TGAAGTGCTTAGCAACTTCTCTGATGTAGATTTCAAGACGCTTGTTCATTACGTCAGGAG  1260
             ||||||||||||||||||||||||||||||||||||||||||||||||||||||||||||
Sbjct  202   TGAAGTGCTTAGCAACTTCTCTGATGTAGATTTCAAGACGCTTGTTCATTACGTCAGGAG  143

Query  1261  TTACATAAGGCTTGGATGTATCATAATCTTCATGGAAGAAGAATGAAGGAGTCTGTG  1317
             |||||||||||||||||||||||||||||||||||||||||||||||||||||||||
Sbjct  142   TTACATAAGGCTTGGATGTATCATAATCTTCATGGAAGAAGAATGAAGGAGTCTGTG  86

Lambda      K        H
    1.33    0.621     1.12

Gapped
Lambda      K        H
    1.28    0.460    0.850

Effective search space used: 208501634

  Database: ../../g2019a.full_hap.fasta
    Posted date:  Jul 24, 2019  2:55 PM
  Number of letters in database: 163,008
  Number of sequences in database:  125

Matrix: blastn matrix 1 -2
Gap Penalties: Existence: 0, Extension: 2.5
```

Listing 1: BLAST output demonstrating 100% alignment of a PacBio CCS read to a haplotype recovered by `Gretel` for G123 (#0).

```
BLASTN 2.7.1+

Reference: Zheng Zhang, Scott Schwartz, Lukas Wagner, and Webb
Miller (2000), "A greedy algorithm for aligning DNA sequences", J
Comput Biol 2000; 7(1-2):203-14.

Database: ../../g2019a.full_hap.fasta
           125 sequences; 163,008 total letters

Query= m54118_190409_113909/70845351/ccs
Length=1317

> G123_2__-27.29
Length=1425

 Score = 2433 bits (1317),  Expect = 0.0
 Identities = 1317/1317 (100%), Gaps = 0/1317 (0%)
 Strand=Plus/Minus

Query  1     CGCTTGCAGATGCCTTAGCACCAACGTTGATAGCTACAGGAAGCTCATCTGCTGTCTTCC  60
             ||||||||||||||||||||||||||||||||||||||||||||||||||||||||||||
Sbjct  1402  CGCTTGCAGATGCCTTAGCACCAACGTTGATAGCTACAGGAAGCTCATCTGCTGTCTTCC  1343

Query  61    AGGCTTCATCATCAAACTCACCATCAATAACAGGTGTGCCCTTCTTAACTACTGCAAGTG  120
             ||||||||||||||||||||||||||||||||||||||||||||||||||||||||||||
Sbjct  1342  AGGCTTCATCATCAAACTCACCATCAATAACAGGTGTGCCCTTCTTAACTACTGCAAGTG  1283

Query  121   GCTTAATTACTGTTTCAGCAAAGAACTTGCTGGACTGAGCATGTTTAAATGTTGTGTCGT  180
             ||||||||||||||||||||||||||||||||||||||||||||||||||||||||||||
Sbjct  1282  GCTTAATTACTGTTTCAGCAAAGAACTTGCTGGACTGAGCATGTTTAAATGTTGTGTCGT  1223

Query  181   TAAATACAGCAAGGAAATCACCGTCATTAACTACAATATCAAACTTAACCTTGTTAGCTG  240
             ||||||||||||||||||||||||||||||||||||||||||||||||||||||||||||
Sbjct  1222  TAAATACAGCAAGGAAATCACCGTCATTAACTACAATATCAAACTTAACCTTGTTAGCTG  1163

Query  241   TAAGAGCTTCGAAGTCAACAGGAACATTTACTACTGCCTCGTATCCATTTTCATTTTCTG  300
             ||||||||||||||||||||||||||||||||||||||||||||||||||||||||||||
Sbjct  1162  TAAGAGCTTCGAAGTCAACAGGAACATTTACTACTGCCTCGTATCCATTTTCATTTTCTG  1103

Query  301   TTGCTTCTTCTCTCTTTACTACTACTGATTTGATTCCTGTTCCGTCATCTGCATAAACTG  360
             ||||||||||||||||||||||||||||||||||||||||||||||||||||||||||||
Sbjct  1102  TTGCTTCTTCTCTCTTTACTACTACTGATTTGATTCCTGTTCCGTCATCTGCATAAACTG  1043

Query  361   TAAAGAAATCATCCTGGCCAGTTGTTGTATCTTTTATTACAACTTTAACGTCGATGCCAT  420
             ||||||||||||||||||||||||||||||||||||||||||||||||||||||||||||
Sbjct  1042  TAAAGAAATCATCCTGGCCAGTTGTTGTATCTTTTATTACAACTTTAACGTCGATGCCAT  983

Query  421   TCTCACTCCACATAGGGATAAACTTGCCAGCAACAAACTCATATACATTACCAGCTGAGA  480
             ||||||||||||||||||||||||||||||||||||||||||||||||||||||||||||
Sbjct  982   TCTCACTCCACATAGGGATAAACTTGCCAGCAACAAACTCATATACATTACCAGCTGAGA  923

Query  481   AATCATTATTTACATTCTGTACAAGTGTTATTGACTTTACTTCAGGCTCAAGTTCTCCGG  540
             ||||||||||||||||||||||||||||||||||||||||||||||||||||||||||||
Sbjct  922   AATCATTATTTACATTCTGTACAAGTGTTATTGACTTTACTTCAGGCTCAAGTTCTCCGG  863

Query  541   CATTTGCAATTGCCCAGAAGCAATCCTTAGCCTTGTAGTAATCATCAAAGAGAAGTGGAT  600
             ||||||||||||||||||||||||||||||||||||||||||||||||||||||||||||
Sbjct  862   CATTTGCAATTGCCCAGAAGCAATCCTTAGCCTTGTAGTAATCATCAAAGAGAAGTGGAT  803

Query  601   ACTGTCTTGCGCTTCCGTCTGCGCCGCCACCGTTATTGTTTGATGACTGGAGCCATGAAT  660
             ||||||||||||||||||||||||||||||||||||||||||||||||||||||||||||
Sbjct  802   ACTGTCTTGCGCTTCCGTCTGCGCCGCCACCGTTATTGTTTGATGACTGGAGCCATGAAT  743

Query  661   GCTTATCTGTTACTCCCCAGATTGTCATACCTGTGAAGTTTACGCCGTCTTCTCTAAGTC  720
             ||||||||||||||||||||||||||||||||||||||||||||||||||||||||||||
Sbjct  742   GCTTATCTGTTACTCCCCAGATTGTCATACCTGTGAAGTTTACGCCGTCTTCTCTAAGTC  683

Query  721   TTCTGATTGTGTCATAAACTGCTTTGTATCTCTCCGCAAGTCTGTCATCCTTGGCACTTG  780
             ||||||||||||||||||||||||||||||||||||||||||||||||||||||||||||
Sbjct  682   TTCTGATTGTGTCATAAACTGCTTTGTATCTCTCCGCAAGTCTGTCATCCTTGGCACTTG  623

Query  781   TAGCACCCTTGAAGTCAAGCTCAGTTACCTGAACCTGGTCAACGATAGCTGCATAAGCCT  840
             ||||||||||||||||||||||||||||||||||||||||||||||||||||||||||||
Sbjct  622   TAGCACCCTTGAAGTCAAGCTCAGTTACCTGAACCTGGTCAACGATAGCTGCATAAGCCT  563

Query  841   TGGCTGCTGTCTTGAACTGCTCCATTGAAGGATTGTTGGATGCAATCTGGTAGTGAGCCT  900
             ||||||||||||||||||||||||||||||||||||||||||||||||||||||||||||
Sbjct  562   TGGCTGCTGTCTTGAACTGCTCCATTGAAGGATTGTTGGATGCAATCTGGTAGTGAGCCT  503

Query  901   GCATACCCATACCATCAATTCTTGCACCAGGTGTTGCCTTAACATCTCTAAGAAGCTGGC  960
             ||||||||||||||||||||||||||||||||||||||||||||||||||||||||||||
Sbjct  502   GCATACCCATACCATCAATTCTTGCACCAGGTGTTGCCTTAACATCTCTAAGAAGCTGGC  443

Query  961   AGATACCACCCATTTTGCTGCTTACTGTCTCGTTGTAATCGTTATAGAAAAGAGCGATAT  1020
             ||||||||||||||||||||||||||||||||||||||||||||||||||||||||||||
Sbjct  442   AGATACCACCCATTTTGCTGCTTACTGTCTCGTTGTAATCGTTATAGAAAAGAGCGATAT  383

Query  1021  CAGCAGGCATATATCTGTTAGCATATACGAATGCATTTGTGATGAACTCCTGTGAGCCGT  1080
             ||||||||||||||||||||||||||||||||||||||||||||||||||||||||||||
Sbjct  382   CAGCAGGCATATATCTGTTAGCATATACGAATGCATTTGTGATGAACTCCTGTGAGCCGT  323

Query  1081  AAACTCCCCACCACTCTGATCTCTCAGATGCATTTCTGTAAGTACCTGTTCCATCACTTA  1140
             ||||||||||||||||||||||||||||||||||||||||||||||||||||||||||||
Sbjct  322   AAACTCCCCACCACTCTGATCTCTCAGATGCATTTCTGTAAGTACCTGTTCCATCACTTA  263

Query  1141  CTGCTTCGTTAACAACATCCCATCCATAGAAGAGATCTTTGTACTTGCTGTCTTCTGCTG  1200
             ||||||||||||||||||||||||||||||||||||||||||||||||||||||||||||
Sbjct  262   CTGCTTCGTTAACAACATCCCATCCATAGAAGAGATCTTTGTACTTGCTGTCTTCTGCTG  203

Query  1201  TGAAGTGCTTAGCAACTTCTCTGATGTAGATTTCAAGACGCTTGTTCATTACGTCAGGAG  1260
             ||||||||||||||||||||||||||||||||||||||||||||||||||||||||||||
Sbjct  202   TGAAGTGCTTAGCAACTTCTCTGATGTAGATTTCAAGACGCTTGTTCATTACGTCAGGAG  143

Query  1261  TTACATAAGGCTTGGATGTATCATAATCTTCATGGAAGAAGAATGAAGGAGTCTGTG  1317
             |||||||||||||||||||||||||||||||||||||||||||||||||||||||||
Sbjct  142   TTACATAAGGCTTGGATGTATCATAATCTTCATGGAAGAAGAATGAAGGAGTCTGTG  86

Lambda      K        H
    1.33    0.621     1.12

Gapped
Lambda      K        H
    1.28    0.460    0.850

Effective search space used: 208501634

  Database: ../../g2019a.full_hap.fasta
    Posted date:  Jul 24, 2019  2:55 PM
  Number of letters in database: 163,008
  Number of sequences in database:  125

Matrix: blastn matrix 1 -2
Gap Penalties: Existence: 0, Extension: 2.5
```

Listing 2: BLAST output demonstrating 100% alignment of a PacBio CCS read to a haplotype recovered by `Gretel` for G123 (#2).

```
BLASTN 2.7.1+

Reference: Zheng Zhang, Scott Schwartz, Lukas Wagner, and Webb
Miller (2000), "A greedy algorithm for aligning DNA sequences", J
Comput Biol 2000; 7(1-2):203-14.

Database: ../../g2019a.full_hap.fasta
           125 sequences; 163,008 total letters

Query= m54118_190409_113909/9634013/ccs
Length=1319

> G123_3__-28.60
Length=1425

 Score = 2423 bits (1312),  Expect = 0.0
 Identities = 1317/1319 (99%), Gaps = 2/1319 (0%)
 Strand=Plus/Minus

Query  1     CGCTTGCAGATGCCTTAGCACCAACGTTGATAGCTACAGGAAGCTCATCTGCTGTCTTCC  60
             ||||||||||||||||||||||||||||||||||||||||||||||||||||||||||||
Sbjct  1402  CGCTTGCAGATGCCTTAGCACCAACGTTGATAGCTACAGGAAGCTCATCTGCTGTCTTCC  1343

Query  61    AGGCTTCATCATCAAACTCACCATCAATAACAGGTGTGCCCTTCTTAACTACTGCAAGTG  120
             ||||||||||||||||||||||||||||||||||||||||||||||||||||||||||||
Sbjct  1342  AGGCTTCATCATCAAACTCACCATCAATAACAGGTGTGCCCTTCTTAACTACTGCAAGTG  1283

Query  121   GCTTAATTACTGTTTCAGCAAAGAACTTGCTGGACTGAGCATGTTTAAATGTTGTGTCGT  180
             ||||||||||||||||||||||||||||||||||||||||||||||||||||||||||||
Sbjct  1282  GCTTAATTACTGTTTCAGCAAAGAACTTGCTGGACTGAGCATGTTTAAATGTTGTGTCGT  1223

Query  181   TAAATACAGCAAGGAAATCACCGTCATTAACTACAATATCAAACTTAACCTTGTTAGCTG  240
             ||||||||||||||||||||||||||||||||||||||||||||||||||||||||||||
Sbjct  1222  TAAATACAGCAAGGAAATCACCGTCATTAACTACAATATCAAACTTAACCTTGTTAGCTG  1163

Query  241   TAAGAGCTTCGAAGTCAACAGGAACATTTACTACTGCCTCGTATCCATTTTCATTTTCTG  300
             ||||||||||||||||||||||||||||||||||||||||||||||||||||||||||||
Sbjct  1162  TAAGAGCTTCGAAGTCAACAGGAACATTTACTACTGCCTCGTATCCATTTTCATTTTCTG  1103

Query  301   TTGCTTCATTCTCTCTTTACTACTACTGATTTGATTCCTGTTCCGTCATCTGCATAAACT  360
             ||||||| ||||||||||||||||||||||||||||||||||||||||||||||||||||
Sbjct  1102  TTGCTTC-TTCTCTCTTTACTACTACTGATTTGATTCCTGTTCCGTCATCTGCATAAACT  1044

Query  361   GTAAAGAAATCATCCTGGCCAGTTGTTGTATCTTTTATTACAACTTTAACGTCGATGCCA  420
             ||||||||||||||||||||||||||||||||||||||||||||||||||||||||||||
Sbjct  1043  GTAAAGAAATCATCCTGGCCAGTTGTTGTATCTTTTATTACAACTTTAACGTCGATGCCA  984

Query  421   TTCTCACTCCACATAGGGATAAACTTGCCAGCAACAAACTCATATACATTACCAGCTGAG  480
             ||||||||||||||||||||||||||||||||||||||||||||||||||||||||||||
Sbjct  983   TTCTCACTCCACATAGGGATAAACTTGCCAGCAACAAACTCATATACATTACCAGCTGAG  924

Query  481   AAATCATTATTTACATTCTGTACAAGTGTTATTGACTTTACTTCAGGCTCAAGTTCTCCG  540
             ||||||||||||||||||||||||||||||||||||||||||||||||||||||||||||
Sbjct  923   AAATCATTATTTACATTCTGTACAAGTGTTATTGACTTTACTTCAGGCTCAAGTTCTCCG  864

Query  541   GCATTTGCAATTGCCCAGAAGCAATCCTTAGCCTTGTAGTAATCATCAAAGAGAAGTGGA  600
             ||||||||||||||||||||||||||||||||||||||||||||||||||||||||||||
Sbjct  863   GCATTTGCAATTGCCCAGAAGCAATCCTTAGCCTTGTAGTAATCATCAAAGAGAAGTGGA  804

Query  601   TACTGTCTTGCGCTTCCGTCTGCAGCCGCCACCGTTATTGTTTGATGACTGGAGCCATGA  660
             ||||||||||||||||||||||| ||||||||||||||||||||||||||||||||||||
Sbjct  803   TACTGTCTTGCGCTTCCGTCTGC-GCCGCCACCGTTATTGTTTGATGACTGGAGCCATGA  745

Query  661   ATGCTTATCTGTTACTCCCCAGATTGTCATACCTGTGAAGTTTACGCCGTCTTCTCTAAG  720
             ||||||||||||||||||||||||||||||||||||||||||||||||||||||||||||
Sbjct  744   ATGCTTATCTGTTACTCCCCAGATTGTCATACCTGTGAAGTTTACGCCGTCTTCTCTAAG  685

Query  721   TCTTCTGATTGTGTCATAAACTGCTTTGTATCTCTCCGCAAGTCTGTCATCCTTGGCACT  780
             ||||||||||||||||||||||||||||||||||||||||||||||||||||||||||||
Sbjct  684   TCTTCTGATTGTGTCATAAACTGCTTTGTATCTCTCCGCAAGTCTGTCATCCTTGGCACT  625

Query  781   TGTAGCACCCTTGAAGTCAAGCTCAGTTACCTGAACCTGGTCAACGATAGCTGCATAAGC  840
             ||||||||||||||||||||||||||||||||||||||||||||||||||||||||||||
Sbjct  624   TGTAGCACCCTTGAAGTCAAGCTCAGTTACCTGAACCTGGTCAACGATAGCTGCATAAGC  565

Query  841   CTTGGCTGCTGTCTTGAACTGCTCCATTGAAGGATTGTTGGATGCAATCTGGTAGTGAGC  900
             ||||||||||||||||||||||||||||||||||||||||||||||||||||||||||||
Sbjct  564   CTTGGCTGCTGTCTTGAACTGCTCCATTGAAGGATTGTTGGATGCAATCTGGTAGTGAGC  505

Query  901   CTGCATACCCATACCATCAATTCTTGCACCAGGTGTTGCCTTAACATCTCTAAGAAGCTG  960
             ||||||||||||||||||||||||||||||||||||||||||||||||||||||||||||
Sbjct  504   CTGCATACCCATACCATCAATTCTTGCACCAGGTGTTGCCTTAACATCTCTAAGAAGCTG  445

Query  961   GCAGATACCACCCATCTTGCTACTTACTGTCTCGTTGTAATCGTTATAGAAAAGAGCAAT  1020
             ||||||||||||||||||||||||||||||||||||||||||||||||||||||||||||
Sbjct  444   GCAGATACCACCCATCTTGCTACTTACTGTCTCGTTGTAATCGTTATAGAAAAGAGCAAT  385

Query  1021  GTCTGCAGGCATATATCTGTTAGCATATACGAATGCATTTGTGATGAACTCCTGTGAGCC  1080
             ||||||||||||||||||||||||||||||||||||||||||||||||||||||||||||
Sbjct  384   GTCTGCAGGCATATATCTGTTAGCATATACGAATGCATTTGTGATGAACTCCTGTGAGCC  325

Query  1081  GTAAACTCCCCACCACTCTGATCTCTCAGATGCATTTCTGTAAGTACCTGTTCCATCACT  1140
             ||||||||||||||||||||||||||||||||||||||||||||||||||||||||||||
Sbjct  324   GTAAACTCCCCACCACTCTGATCTCTCAGATGCATTTCTGTAAGTACCTGTTCCATCACT  265

Query  1141  TACTGCTTCGTTAACAACATCCCATCCATAGAAGAGATCTTTGTACTTGCTGTCTTCTGC  1200
             ||||||||||||||||||||||||||||||||||||||||||||||||||||||||||||
Sbjct  264   TACTGCTTCGTTAACAACATCCCATCCATAGAAGAGATCTTTGTACTTGCTGTCTTCTGC  205

Query  1201  TGTGAAGTGCTTAGCAACTTCTCTGATGTAGATTTCAAGACGCTTGTTCATTACGTCAGG  1260
             ||||||||||||||||||||||||||||||||||||||||||||||||||||||||||||
Sbjct  204   TGTGAAGTGCTTAGCAACTTCTCTGATGTAGATTTCAAGACGCTTGTTCATTACGTCAGG  145

Query  1261  AGTTACATAAGGCTTGGATGTATCATAATCTTCATGGAAGAAGAATGAAGGAGTCTGTG  1319
             |||||||||||||||||||||||||||||||||||||||||||||||||||||||||||
Sbjct  144   AGTTACATAAGGCTTGGATGTATCATAATCTTCATGGAAGAAGAATGAAGGAGTCTGTG  86

Lambda      K        H
    1.33    0.621     1.12

Gapped
Lambda      K        H
    1.28    0.460    0.850

Effective search space used: 208822900

  Database: ../../g2019a.full_hap.fasta
    Posted date:  Jul 24, 2019  2:55 PM
  Number of letters in database: 163,008
  Number of sequences in database:  125

Matrix: blastn matrix 1 -2
Gap Penalties: Existence: 0, Extension: 2.5
```

Listing 3: BLAST output demonstrating 100% alignment (other than 2 disregarded insertions) of a PacBio CCS read to a haplotype recovered by `Gretel` for G123 (#3).

```
BLASTN 2.7.1+

Reference: Zheng Zhang, Scott Schwartz, Lukas Wagner, and Webb
Miller (2000), "A greedy algorithm for aligning DNA sequences", J
Comput Biol 2000; 7(1-2):203-14.

Database: ../../g2019a.full_hap.fasta
           125 sequences; 163,008 total letters

Query= m54118_190409_113909/74580608/ccs
Length=1317

> G123_5__-32.79
Length=1425

 Score = 2433 bits (1317),  Expect = 0.0
 Identities = 1317/1317 (100%), Gaps = 0/1317 (0%)
 Strand=Plus/Minus

Query  1     CGCTTGCAGATGCCTTAGCACCAACGTTGATAGCTACAGGAAGCTCATCTGCTGTCTTCC  60
             ||||||||||||||||||||||||||||||||||||||||||||||||||||||||||||
Sbjct  1402  CGCTTGCAGATGCCTTAGCACCAACGTTGATAGCTACAGGAAGCTCATCTGCTGTCTTCC  1343

Query  61    AGGCTTCATCATCAAACTCACCATCAATAACAGGTGTGCCCTTCTTAACTACTGCAAGTG  120
             ||||||||||||||||||||||||||||||||||||||||||||||||||||||||||||
Sbjct  1342  AGGCTTCATCATCAAACTCACCATCAATAACAGGTGTGCCCTTCTTAACTACTGCAAGTG  1283

Query  121   GCTTAATTACTGTTTCAGCAAAGAACTTGCTGGACTGAGCATGCTTAAATGTTGTGTCGT  180
             ||||||||||||||||||||||||||||||||||||||||||||||||||||||||||||
Sbjct  1282  GCTTAATTACTGTTTCAGCAAAGAACTTGCTGGACTGAGCATGCTTAAATGTTGTGTCGT  1223

Query  181   TAAATACAGCAAGGAAATCACCGTCATTAACTACAATATCAAACTTAACCTTGTTAGCTG  240
             ||||||||||||||||||||||||||||||||||||||||||||||||||||||||||||
Sbjct  1222  TAAATACAGCAAGGAAATCACCGTCATTAACTACAATATCAAACTTAACCTTGTTAGCTG  1163

Query  241   TAAGAGCTTCGAAGTCAACAGGAACATTTACTACTGCCTCGTATCCATTTTCATTTTCTG  300
             ||||||||||||||||||||||||||||||||||||||||||||||||||||||||||||
Sbjct  1162  TAAGAGCTTCGAAGTCAACAGGAACATTTACTACTGCCTCGTATCCATTTTCATTTTCTG  1103

Query  301   TTGCTTCTTCTCTCTTTACTACTACTGATTTGATTCCTGTTCCGTCATCTGCATAAACTG  360
             ||||||||||||||||||||||||||||||||||||||||||||||||||||||||||||
Sbjct  1102  TTGCTTCTTCTCTCTTTACTACTACTGATTTGATTCCTGTTCCGTCATCTGCATAAACTG  1043

Query  361   TAAAGAAATCATCCTGGCCAGTTGTTGTATCTTTTATTACAACTTTAACGTCGATGCCAT  420
             ||||||||||||||||||||||||||||||||||||||||||||||||||||||||||||
Sbjct  1042  TAAAGAAATCATCCTGGCCAGTTGTTGTATCTTTTATTACAACTTTAACGTCGATGCCAT  983

Query  421   TCTCACTCCACATAGGGATAAACTTGCCAGCAACAAACTCATATACATTACCAGCTGAGA  480
             ||||||||||||||||||||||||||||||||||||||||||||||||||||||||||||
Sbjct  982   TCTCACTCCACATAGGGATAAACTTGCCAGCAACAAACTCATATACATTACCAGCTGAGA  923

Query  481   AATCATTATTTACATTCTGTACAAGTGTTATTGACTTTACTTCAGGCTCAAGTTCTCCGG  540
             ||||||||||||||||||||||||||||||||||||||||||||||||||||||||||||
Sbjct  922   AATCATTATTTACATTCTGTACAAGTGTTATTGACTTTACTTCAGGCTCAAGTTCTCCGG  863

Query  541   CATTTGCAATTGCCCAGAAGCAATCCTTAGCCTTGTAGTAATCATCAAAGAGAAGTGGAT  600
             ||||||||||||||||||||||||||||||||||||||||||||||||||||||||||||
Sbjct  862   CATTTGCAATTGCCCAGAAGCAATCCTTAGCCTTGTAGTAATCATCAAAGAGAAGTGGAT  803

Query  601   ACTGTCTTGCGCTTCCGTCTGCGCCGCCACCGTTATTGTTTGATGACTGGAGCCATGAAT  660
             ||||||||||||||||||||||||||||||||||||||||||||||||||||||||||||
Sbjct  802   ACTGTCTTGCGCTTCCGTCTGCGCCGCCACCGTTATTGTTTGATGACTGGAGCCATGAAT  743

Query  661   GCTTATCTGTTACTCCCCAGATTGTCATACCTGTGAAGTTTACGCCGTCTTCTCTAAGTC  720
             ||||||||||||||||||||||||||||||||||||||||||||||||||||||||||||
Sbjct  742   GCTTATCTGTTACTCCCCAGATTGTCATACCTGTGAAGTTTACGCCGTCTTCTCTAAGTC  683

Query  721   TTCTGATTGTGTCATAAACTGCTTTGTATCTCTCCGCAAGTCTGTCATCCTTGGCACTTG  780
             ||||||||||||||||||||||||||||||||||||||||||||||||||||||||||||
Sbjct  682   TTCTGATTGTGTCATAAACTGCTTTGTATCTCTCCGCAAGTCTGTCATCCTTGGCACTTG  623

Query  781   TAGCACCCTTGAAGTCAAGCTCAGTTACCTGAACCTGGTCAACGATAGCTGCATAAGCCT  840
             ||||||||||||||||||||||||||||||||||||||||||||||||||||||||||||
Sbjct  622   TAGCACCCTTGAAGTCAAGCTCAGTTACCTGAACCTGGTCAACGATAGCTGCATAAGCCT  563

Query  841   TGGCTGCTGTCTTGAACTGCTCCATTGAAGGATTGTTGGATGCAATCTGGTAGTGAGCCT  900
             ||||||||||||||||||||||||||||||||||||||||||||||||||||||||||||
Sbjct  562   TGGCTGCTGTCTTGAACTGCTCCATTGAAGGATTGTTGGATGCAATCTGGTAGTGAGCCT  503

Query  901   GCATACCCATACCATCAATTCTTGCACCAGGTGTTGCCTTAACATCTCTAAGAAGCTGGC  960
             ||||||||||||||||||||||||||||||||||||||||||||||||||||||||||||
Sbjct  502   GCATACCCATACCATCAATTCTTGCACCAGGTGTTGCCTTAACATCTCTAAGAAGCTGGC  443

Query  961   AGATACCACCCATTTTGCTGCTTACTGTCTCGTTGTAATCGTTATAGAAAAGAGCGATAT  1020
             ||||||||||||||||||||||||||||||||||||||||||||||||||||||||||||
Sbjct  442   AGATACCACCCATTTTGCTGCTTACTGTCTCGTTGTAATCGTTATAGAAAAGAGCGATAT  383

Query  1021  CAGCAGGCATATATCTGTTAGCATATACGAATGCATTTGTGATGAACTCCTGTGAGCCGT  1080
             ||||||||||||||||||||||||||||||||||||||||||||||||||||||||||||
Sbjct  382   CAGCAGGCATATATCTGTTAGCATATACGAATGCATTTGTGATGAACTCCTGTGAGCCGT  323

Query  1081  AAACTCCCCACCACTCTGATCTCTCAGATGCATTTCTGTAAGTACCTGTTCCATCACTTA  1140
             ||||||||||||||||||||||||||||||||||||||||||||||||||||||||||||
Sbjct  322   AAACTCCCCACCACTCTGATCTCTCAGATGCATTTCTGTAAGTACCTGTTCCATCACTTA  263

Query  1141  CTGCTTCGTTAACAACATCCCATCCATAGAAGAGATCTTTGTACTTGCTGTCTTCTGCTG  1200
             ||||||||||||||||||||||||||||||||||||||||||||||||||||||||||||
Sbjct  262   CTGCTTCGTTAACAACATCCCATCCATAGAAGAGATCTTTGTACTTGCTGTCTTCTGCTG  203

Query  1201  TGAAGTGCTTAGCAACTTCTCTGATGTAGATTTCAAGACGCTTGTTCATTACGTCAGGAG  1260
             ||||||||||||||||||||||||||||||||||||||||||||||||||||||||||||
Sbjct  202   TGAAGTGCTTAGCAACTTCTCTGATGTAGATTTCAAGACGCTTGTTCATTACGTCAGGAG  143

Query  1261  TTACATAAGGCTTGGATGTATCATAATCTTCATGGAAGAAGAATGAAGGAGTCTGTG  1317
             |||||||||||||||||||||||||||||||||||||||||||||||||||||||||
Sbjct  142   TTACATAAGGCTTGGATGTATCATAATCTTCATGGAAGAAGAATGAAGGAGTCTGTG  86

Lambda      K        H
    1.33    0.621     1.12

Gapped
Lambda      K        H
    1.28    0.460    0.850

Effective search space used: 208501634

  Database: ../../g2019a.full_hap.fasta
    Posted date:  Jul 24, 2019  2:55 PM
  Number of letters in database: 163,008
  Number of sequences in database:  125

Matrix: blastn matrix 1 -2
Gap Penalties: Existence: 0, Extension: 2.5
```

Listing 4: BLAST output demonstrating 100% alignment of a PacBio CCS read to a haplotype recovered by `Gretel` for G123 (#5).

```
BLASTN 2.7.1+

Reference: Zheng Zhang, Scott Schwartz, Lukas Wagner, and Webb
Miller (2000), "A greedy algorithm for aligning DNA sequences", J
Comput Biol 2000; 7(1-2):203-14.

Database: ../../g2019a.full_hap.fasta
           125 sequences; 163,008 total letters

Query= m54118_190409_113909/7537133/ccs
Length=1317

> G123_8__-36.02
Length=1425

 Score = 2433 bits (1317),  Expect = 0.0
 Identities = 1317/1317 (100%), Gaps = 0/1317 (0%)
 Strand=Plus/Plus

Query  1     CACAGACTCCTTCATTCTTCTTCCATGAAGATTATGATACATCCAAGCCTTATGTAACTC  60
             ||||||||||||||||||||||||||||||||||||||||||||||||||||||||||||
Sbjct  86    CACAGACTCCTTCATTCTTCTTCCATGAAGATTATGATACATCCAAGCCTTATGTAACTC  145

Query  61    CTGACGTAATGAACAAGCGTCTTGAAATCTACATCAGAGAAGTTGCTAAGCACTTCACAG  120
             ||||||||||||||||||||||||||||||||||||||||||||||||||||||||||||
Sbjct  146   CTGACGTAATGAACAAGCGTCTTGAAATCTACATCAGAGAAGTTGCTAAGCACTTCACAG  205

Query  121   CAGAAGACAGCAAGTACAAAGATCTCTTCTATGGATGGGATGTTGTTAACGAAGCAGTAA  180
             ||||||||||||||||||||||||||||||||||||||||||||||||||||||||||||
Sbjct  206   CAGAAGACAGCAAGTACAAAGATCTCTTCTATGGATGGGATGTTGTTAACGAAGCAGTAA  265

Query  181   GTGATGGAACAGGTACTTACAGAAATGCATCTGAGAGATCAGAGTGGTGGGGAGTTTACG  240
             ||||||||||||||||||||||||||||||||||||||||||||||||||||||||||||
Sbjct  266   GTGATGGAACAGGTACTTACAGAAATGCATCTGAGAGATCAGAGTGGTGGGGAGTTTACG  325

Query  241   GCTCACAGGAGTTCATCACAAATGCATTCGTATATGCTAACAGATATATGCCTGCTGATA  300
             ||||||||||||||||||||||||||||||||||||||||||||||||||||||||||||
Sbjct  326   GCTCACAGGAGTTCATCACAAATGCATTCGTATATGCTAACAGATATATGCCTGCTGATA  385

Query  301   TCGCTCTTTTCTATAACGATTATAACGAGACAGTAAGCAGCAAGATGGGTGGTATCTGCC  360
             ||||||||||||||||||||||||||||||||||||||||||||||||||||||||||||
Sbjct  386   TCGCTCTTTTCTATAACGATTATAACGAGACAGTAAGCAGCAAGATGGGTGGTATCTGCC  445

Query  361   AGCTTCTTAGAGATGTTAAGGCAACACCTGGTGCAAGAATTGATGGTATGGGTATGCAGG  420
             ||||||||||||||||||||||||||||||||||||||||||||||||||||||||||||
Sbjct  446   AGCTTCTTAGAGATGTTAAGGCAACACCTGGTGCAAGAATTGATGGTATGGGTATGCAGG  505

Query  421   CTCACTACCAGATTGCATCCAACAATCCTTCAATGGAGCAGTTCAAGACAGCAGCCAAGG  480
             ||||||||||||||||||||||||||||||||||||||||||||||||||||||||||||
Sbjct  506   CTCACTACCAGATTGCATCCAACAATCCTTCAATGGAGCAGTTCAAGACAGCAGCCAAGG  565

Query  481   CTTATGCAGCTATCGTTGACCAGGTTCAGGTAACTGAGCTTGACTTCAAGGGTGCTACAA  540
             ||||||||||||||||||||||||||||||||||||||||||||||||||||||||||||
Sbjct  566   CTTATGCAGCTATCGTTGACCAGGTTCAGGTAACTGAGCTTGACTTCAAGGGTGCTACAA  625

Query  541   GTGCCAAGGATGACAGACTTGCGGAGAGATACAAAGCAGTTTATGACACAATCAGAAGAC  600
             ||||||||||||||||||||||||||||||||||||||||||||||||||||||||||||
Sbjct  626   GTGCCAAGGATGACAGACTTGCGGAGAGATACAAAGCAGTTTATGACACAATCAGAAGAC  685

Query  601   TTAGAGAAGACGGCGTAAACTTCACAGGTATGACAATCTGGGGAGTAACAGATAAGCATT  660
             ||||||||||||||||||||||||||||||||||||||||||||||||||||||||||||
Sbjct  686   TTAGAGAAGACGGCGTAAACTTCACAGGTATGACAATCTGGGGAGTAACAGATAAGCATT  745

Query  661   CATGGCTCCAGTCATCAAACAATAACGGTGGCGGCGCAGACGGAAGCGCAAGACAGTATC  720
             ||||||||||||||||||||||||||||||||||||||||||||||||||||||||||||
Sbjct  746   CATGGCTCCAGTCATCAAACAATAACGGTGGCGGCGCAGACGGAAGCGCAAGACAGTATC  805

Query  721   CACTTCTCTTTGATGATTACTACAAGGCTAAGGATTGCTTCTGGGCAATTGCAAATGCCG  780
             ||||||||||||||||||||||||||||||||||||||||||||||||||||||||||||
Sbjct  806   CACTTCTCTTTGATGATTACTACAAGGCTAAGGATTGCTTCTGGGCAATTGCAAATGCCG  865

Query  781   GAGAACTTGAGCCTGAAGTAAAGTCAATAACACTTGTACAGAATGTAAATAATGATTTCT  840
             ||||||||||||||||||||||||||||||||||||||||||||||||||||||||||||
Sbjct  866   GAGAACTTGAGCCTGAAGTAAAGTCAATAACACTTGTACAGAATGTAAATAATGATTTCT  925

Query  841   CAGCTGGTAATGTATATGAGTTTGTTGCTGGCAAGTTTATCCCTATGTGGAGTGAGAATG  900
             ||||||||||||||||||||||||||||||||||||||||||||||||||||||||||||
Sbjct  926   CAGCTGGTAATGTATATGAGTTTGTTGCTGGCAAGTTTATCCCTATGTGGAGTGAGAATG  985

Query  901   GCATCGACGTTAAAGTTGTAATAAAAGATACAACAACTGGCCAGGATGATTTCTTTACAG  960
             ||||||||||||||||||||||||||||||||||||||||||||||||||||||||||||
Sbjct  986   GCATCGACGTTAAAGTTGTAATAAAAGATACAACAACTGGCCAGGATGATTTCTTTACAG  1045

Query  961   TTTATGCAGATGACGGAACAGGAATCAAATCAGTAGTAGTAAAGAGAGAAGAAGCAACAG  1020
             ||||||||||||||||||||||||||||||||||||||||||||||||||||||||||||
Sbjct  1046  TTTATGCAGATGACGGAACAGGAATCAAATCAGTAGTAGTAAAGAGAGAAGAAGCAACAG  1105

Query  1021  AAAATGAAAATGGATACGAGGCAGTAGTAAATGTTCCTGTTGACTTCGAAGCTCTTACAG  1080
             ||||||||||||||||||||||||||||||||||||||||||||||||||||||||||||
Sbjct  1106  AAAATGAAAATGGATACGAGGCAGTAGTAAATGTTCCTGTTGACTTCGAAGCTCTTACAG  1165

Query  1081  CTAACAAGGTTAAGTTTGATATTGTAGTTAATGACGGTGATTTCCTTGCTGTATTTAACG  1140
             ||||||||||||||||||||||||||||||||||||||||||||||||||||||||||||
Sbjct  1166  CTAACAAGGTTAAGTTTGATATTGTAGTTAATGACGGTGATTTCCTTGCTGTATTTAACG  1225

Query  1141  ACACAACATTTAAACATGCTCAGTCCAGCAAGTTCTTTGCTGAAACAGTAATTAAGCCAC  1200
             ||||||||||||||||||||||||||||||||||||||||||||||||||||||||||||
Sbjct  1226  ACACAACATTTAAACATGCTCAGTCCAGCAAGTTCTTTGCTGAAACAGTAATTAAGCCAC  1285

Query  1201  TTGCAGTAGTTAAGAAGGGCACACCTGTTATTGATGGTGAGTTTGATGATGAAGCCTGGA  1260
             ||||||||||||||||||||||||||||||||||||||||||||||||||||||||||||
Sbjct  1286  TTGCAGTAGTTAAGAAGGGCACACCTGTTATTGATGGTGAGTTTGATGATGAAGCCTGGA  1345

Query  1261  AGACAGCAGATGAGCTTCCTGTAGCTATCAACGTTGGTGCTAAGGCATCTGCAAGCG  1317
             |||||||||||||||||||||||||||||||||||||||||||||||||||||||||
Sbjct  1346  AGACAGCAGATGAGCTTCCTGTAGCTATCAACGTTGGTGCTAAGGCATCTGCAAGCG  1402

Lambda      K        H
    1.33    0.621     1.12

Gapped
Lambda      K        H
    1.28    0.460    0.850

Effective search space used: 208501634

  Database: ../../g2019a.full_hap.fasta
    Posted date:  Jul 24, 2019  2:55 PM
  Number of letters in database: 163,008
  Number of sequences in database:  125

Matrix: blastn matrix 1 -2
Gap Penalties: Existence: 0, Extension: 2.5
```

Listing 5: BLAST output demonstrating 100% alignment of a PacBio CCS read to a haplotype recovered by `Gretel` for G123 (#8).

#### 11.4 CCS reads perfectly match G90 haplotypes

```
BLASTN 2.7.1+

Reference: Zheng Zhang, Scott Schwartz, Lukas Wagner, and Webb
Miller (2000), "A greedy algorithm for aligning DNA sequences", J
Comput Biol 2000; 7(1-2):203-14.

Database: ../../g2019a.full_hap.fasta
           125 sequences; 163,008 total letters

Query= m54118_190409_113909/34734233/ccs
Length=812

> G90_0__-28.13
Length=921

 Score = 1493 bits (808),  Expect = 0.0
 Identities = 811/812 (99%), Gaps = 1/812 (0%)
 Strand=Plus/Plus

Query  1    CAATCGACTTAGCTGGCACCTTCATGGTAAGCACGCCCTTCTTAATCTTAGCGTCCTTGT  60
            ||||||||||||||||||||||||||||||||||||||||||||||||||||||||||||
Sbjct  19   CAATCGACTTAGCTGGCACCTTCATGGTAAGCACGCCCTTCTTAATCTTAGCGTCCTTGT  78

Query  61   ACTCCTTTGGAGCAACCACGTTAGGATGATTAAAGTCGTTGTAGTCGGCTACATTCTTTG  120
            ||||||||||||||||||||||||||||||||||||||||||||||||||||||||||||
Sbjct  79   ACTCCTTTGGAGCAACCACGTTAGGATGATTAAAGTCGTTGTAGTCGGCTACATTCTTTG  138

Query  121  AGGTGAGGATGCGGCCGCTAACCTGCTTGGCGTTGAAACCAGCGAAGTTGAAATCTACGG  180
            ||||||||||||||||||||||||||||||||||||||||||||||||||||||||||||
Sbjct  139  AGGTGAGGATGCGGCCGCTAACCTGCTTGGCGTTGAAACCAGCGAAGTTGAAATCTACGG  198

Query  181  TCTGATCCTTATCCAGACTTACGTTTGTGATGGCCAGAACGATGCTGCCATCCTTGGTCT  240
            ||||||||||||||||||||||||||||||||||||||||||||||||||||||||||||
Sbjct  199  TCTGATCCTTATCCAGACTTACGTTTGTGATGGCCAGAACGATGCTGCCATCCTTGGTCT  258

Query  241  TAGCTGCCGATGCAGAGAGCAGTGGGCAAGGACGTGTGGTGTTGTCGCTGGTCTTCTGCT  300
            ||||||||||||||||||||||||||||||||||||||||||||||||||||||||||||
Sbjct  259  TAGCTGCCGATGCAGAGAGCAGTGGGCAAGGACGTGTGGTGTTGTCGCTGGTCTTCTGCT  318

Query  301  TCTCCTTCCAGTACTCGGTGCTGCACTGCACAGTCTCGCTCTGCAGATCAACAGGCAGAT  360
            ||||||||||||||||||||||||||||||||||||||||||||||||||||||||||||
Sbjct  319  TCTCCTTCCAGTACTCGGTGCTGCACTGCACAGTCTCGCTCTGCAGATCAACAGGCAGAT  378

Query  361  AAGTAGCCTCCTGGAATGGGGTGTACATCTGGAATACGTGATAGGTTGGGGTGAGAACCA  420
            ||||||||||||||||||||||||||||||||||||||||||||||||||||||||||||
Sbjct  379  AAGTAGCCTCCTGGAATGGGGTGTACATCTGGAATACGTGATAGGTTGGGGTGAGAACCA  438

Query  421  TGTGGCCTGTGCCCTCCTGATCAGTCAGAATCATCGACTGCAGCACGTTAACCACCTGAG  480
            ||||||||||||||||||||||||||||||||||||||||||||||||||||||||||||
Sbjct  439  TGTGGCCTGTGCCCTCCTGATCAGTCAGAATCATCGACTGCAGCACGTTAACCACCTGAG  498

Query  481  CGATGTTAGCCATCTTTACACGGTCGGTGTACTTATGGAACACATTCAGACTCAGAGCTG  540
            ||||||||||||||||||||||||||||||||||||||||||||||||||||||||||||
Sbjct  499  CGATGTTAGCCATCTTTACACGGTCGGTGTACTTATGGAACACATTCAGACTCAGAGCTG  558

Query  541  CTACGAAGGCATCGCGCAGGGCGTTCTGCTGGTACAGGTGGCCCAGGGATAGTGCCGGGC  600
            ||||||||||||||||||||||||||||||||||||||||| ||||||||||||||||||
Sbjct  559  CTACGAAGGCATCGCGCAGGGCGTTCTGCTGGTACAGGTGG-CCAGGGATAGTGCCGGGC  617

Query  601  TCCTCGTCCCACCAGGTTCCCCACTCATCAACCAGCAGGCCGATGTTGTTCTGAGGATCC  660
            ||||||||||||||||||||||||||||||||||||||||||||||||||||||||||||
Sbjct  618  TCCTCGTCCCACCAGGTTCCCCACTCATCAACCAGCAGGCCGATGTTGTTCTGAGGATCC  677

Query  661  TTCTCGTCCATGATGGCCTTGTGCTTCTTGATTACCTCCTCGATCTCCAAGCACTTACCC  720
            ||||||||||||||||||||||||||||||||||||||||||||||||||||||||||||
Sbjct  678  TTCTCGTCCATGATGGCCTTGTGCTTCTTGATTACCTCCTCGATCTCCAAGCACTTACCC  737

Query  721  AGTGCCCAGTAGTACTGATCGTTATCGAAGTTGATGGCCGAACCCTTAGGACCATTCCAA  780
            ||||||||||||||||||||||||||||||||||||||||||||||||||||||||||||
Sbjct  738  AGTGCCCAGTAGTACTGATCGTTATCGAAGTTGATGGCCGAACCCTTAGGACCATTCCAA  797

Query  781  CCGGTGCAGGTGTAGTAGTGCAGAGAGATACC  812
            ||||||||||||||||||||||||||||||||
Sbjct  798  CCGGTGCAGGTGTAGTAGTGCAGAGAGATACC  829

Lambda      K        H
    1.33    0.621     1.12

Gapped
Lambda      K        H
    1.28    0.460    0.850

Effective search space used: 127641852

  Database: ../../g2019a.full_hap.fasta
    Posted date:  Jul 24, 2019  2:55 PM
  Number of letters in database: 163,008
  Number of sequences in database:  125

Matrix: blastn matrix 1 -2
Gap Penalties: Existence: 0, Extension: 2.5
```

Listing 6: BLAST output demonstrating 100% alignment (other than 1 disregarded insertion) of a PacBio CCS read to a haplotype recovered by `Gretel` for G90 (#0).

```
BLASTN 2.7.1+

Reference: Zheng Zhang, Scott Schwartz, Lukas Wagner, and Webb
Miller (2000), "A greedy algorithm for aligning DNA sequences", J
Comput Biol 2000; 7(1-2):203-14.

Database: ../../g2019a.full_hap.fasta
           125 sequences; 163,008 total letters

Query= m54118_190409_113909/65864248/ccs
Length=814

> G90_1__-50.64
Length=921

 Score = 1483 bits (803),  Expect = 0.0
 Identities = 811/814 (99%), Gaps = 3/814 (0%)
 Strand=Plus/Plus

Query  1    CAATCGACTTAGCTGGCACCTTCATGGTAAGCACGCCCTTCTTAATCTTAGCGTCCTTGT  60
            ||||||||||||||||||||||||||||||||||||||||||||||||||||||||||||
Sbjct  19   CAATCGACTTAGCTGGCACCTTCATGGTAAGCACGCCCTTCTTAATCTTAGCGTCCTTGT  78

Query  61   ACTCCTTTGGAGCTACCACGTTAGGATGATTAAAGTCGTTGTAGTCGGCTACATTCTTTG  120
            ||||||||||||||||||||||||||||||||||||||||||||||||||||||||||||
Sbjct  79   ACTCCTTTGGAGCTACCACGTTAGGATGATTAAAGTCGTTGTAGTCGGCTACATTCTTTG  138

Query  121  AGGTGAGGATGCGGCCGCTAACCTGCTTGGCATTGAAACCAGCGAAGTTGAAATCTACGG  180
            ||||||||||||||||||||||||||||||||||||||||||||||||||||||||||||
Sbjct  139  AGGTGAGGATGCGGCCGCTAACCTGCTTGGCATTGAAACCAGCGAAGTTGAAATCTACGG  198

Query  181  TCTGATCCTTATCCAGACTTACGTTTGTGATGGCCAGAACGATGCTGCCATCCTTGGTCT  240
            ||||||||||||||||||||||||||||||||||||||||||||||||||||||||||||
Sbjct  199  TCTGATCCTTATCCAGACTTACGTTTGTGATGGCCAGAACGATGCTGCCATCCTTGGTCT  258

Query  241  TAGCTGCCGATGCAGAGAGCAGTGGGCAAGGTACGTGTGGTGTTGTCGCTGGTCTTCTGC  300
            ||||||||||||||||||||||||||||||| ||||||||||||||||||||||||||||
Sbjct  259  TAGCTGCCGATGCAGAGAGCAGTGGGCAAGG-ACGTGTGGTGTTGTCGCTGGTCTTCTGC  317

Query  301  TTTTCCTTCCAGTACTCGGTGCTGCACTGCACAGTCTCGCTCTGCAGATCAACAGGCAGA  360
            ||||||||||||||||||||||||||||||||||||||||||||||||||||||||||||
Sbjct  318  TTTTCCTTCCAGTACTCGGTGCTGCACTGCACAGTCTCGCTCTGCAGATCAACAGGCAGA  377

Query  361  TAAGTAGCCTCCTGGAATGGGGTGTACATCTGGAACTACGTGATAGGTTGGGGTGAGAAC  420
            |||||||||||||||||||||||||||||||||||| |||||||||||||||||||||||
Sbjct  378  TAAGTAGCCTCCTGGAATGGGGTGTACATCTGGAAC-ACGTGATAGGTTGGGGTGAGAAC  436

Query  421  CATGTGGCCTGTGCCCTCCTGATCAGTCAGAATCATCGACTGCAGCACGTTAACCACCTG  480
            ||||||||||||||||||||||||||||||||||||||||||||||||||||||||||||
Sbjct  437  CATGTGGCCTGTGCCCTCCTGATCAGTCAGAATCATCGACTGCAGCACGTTAACCACCTG  496

Query  481  AGCGATGTTAGCCATCTTTACACGGTCGGTGTACTTATGGAACACATTCAGACTCAGAGC  540
            ||||||||||||||||||||||||||||||||||||||||||||||||||||||||||||
Sbjct  497  AGCGATGTTAGCCATCTTTACACGGTCGGTGTACTTATGGAACACATTCAGACTCAGAGC  556

Query  541  TGCTACGAAGGCATCGCGCAGGGCGTTCTGCTGGTACAGGTGGCCCAGGGATAGTGCCGG  600
            ||||||||||||||||||||||||||||||||||||||||||| ||||||||||||||||
Sbjct  557  TGCTACGAAGGCATCGCGCAGGGCGTTCTGCTGGTACAGGTGG-CCAGGGATAGTGCCGG  615

Query  601  GCTCCTCGTCCCACCAGGTTCCCCACTCATCAACCAGCAGGCCGATGTTGTTCTGAGGAT  660
            ||||||||||||||||||||||||||||||||||||||||||||||||||||||||||||
Sbjct  616  GCTCCTCGTCCCACCAGGTTCCCCACTCATCAACCAGCAGGCCGATGTTGTTCTGAGGAT  675

Query  661  CCTTCTCGTCCATGATGGCCTTGTGCTTCTTGATTACCTCCTCGATCTCCAAGCACTTAC  720
            ||||||||||||||||||||||||||||||||||||||||||||||||||||||||||||
Sbjct  676  CCTTCTCGTCCATGATGGCCTTGTGCTTCTTGATTACCTCCTCGATCTCCAAGCACTTAC  735

Query  721  CCAGTGCCCAGTAGTACTGATCGTTATCGAAGTTGATGGCCGAACCCTTAGGACCATTCC  780
            ||||||||||||||||||||||||||||||||||||||||||||||||||||||||||||
Sbjct  736  CCAGTGCCCAGTAGTACTGATCGTTATCGAAGTTGATGGCCGAACCCTTAGGACCATTCC  795

Query  781  AACCGGTGCAGGTGTAGTAGTGCAGAGAGATACC  814
            ||||||||||||||||||||||||||||||||||
Sbjct  796  AACCGGTGCAGGTGTAGTAGTGCAGAGAGATACC  829

Lambda      K        H
    1.33    0.621     1.12

Gapped
Lambda      K        H
    1.28    0.460    0.850

Effective search space used: 127963368

  Database: ../../g2019a.full_hap.fasta
    Posted date:  Jul 24, 2019  2:55 PM
  Number of letters in database: 163,008
  Number of sequences in database:  125

Matrix: blastn matrix 1 -2
Gap Penalties: Existence: 0, Extension: 2.5
```

Listing 7: BLAST output demonstrating 100% alignment (other than 3 disregarded insertions) of a PacBio CCS read to a haplotype recovered by `Gretel` for G90 (#1).

#### 11.5 G31 CCS reads do not support Gretel haplotypes

| Haplotype | #CCS | #Mismatches | Mismatch SNP Pos. | Representative Read | #Subreads |
|---|---|---|---|---|---|
| G31_1 | 12 | 19 | 77, 93, 107, 113, 114, 128, 131, 132, 154, 156, 171, 192, 196, 219, 227, 250, 251, 430, 433 | m54118_190409_113909/19202849/ccs | 62 |
| G31_1 | 16 | 50 | 55, 69, 70, 93, 131, 132, 148, 162, 166, 169, 228, 234, 239, 242, 251, 257, 265, 284, 295, 311, 356, 360, 361, 362, 363, 365, 367, 372, 373, 374, 385, 386, 387, 391, 393, 400, 401, 419, 427, 430, 432, 435, 436, 444, 445, 450, 452, 454, 457, 460 | m54118_190409_113909/61276392/ccs | 70 |
| G31_1 | 18 | 18 | 37, 77, 93, 107, 113, 114, 128, 131, 132, 154, 156, 171, 192, 196, 219, 227, 250, 251 | m54118_190409_113909/74252629/ccs | 68 |
| G31_1 | 19 | 39 | 63, 69, 70, 93, 107, 113, 114, 134, 162, 166, 228, 234, 251, 257, 265, 284, 295, 354, 360, 361, 362, 363, 365, 367, 372, 373, 374, 385, 386, 387, 391, 393, 400, 401, 408, 411, 430, 435, 436 | m54118_190409_113909/26345547/ccs | 59 |
| G31_2 | 25 | 20 | 141, 147, 176, 184, 185, 192, 193, 196, 206, 229, 230, 232, 234, 235, 236, 237, 238, 239, 401, 430 | m54118_190409_113909/5243853/ccs | 43 |
| G31_1 | 31 | 19 | 37, 77, 93, 107, 113, 114, 128, 131, 132, 154, 156, 171, 192, 196, 219, 227, 250, 251, 430 | m54118_190409_113909/5570807/ccs | 73 |
| G31_0 | 33 | 50 | 69, 70, 127, 131, 132, 148, 166, 169, 185, 228, 229, 234, 235, 236, 237, 238, 239, 242, 284, 311, 356, 360, 361, 362, 363, 365, 367, 372, 373, 374, 385, 386, 387, 391, 393, 400, 401, 419, 427, 430, 432, 435, 436, 444, 445, 450, 452, 454, 457, 460 | m54118_190409_113909/71630982/ccs | 65 |
| G31_0 | 36 | 15 | 120, 141, 147, 176, 185, 199, 200, 201, 230, 232, 239, 251, 257, 265, 295 | m54118_190409_113909/64159821/ccs | 78 |
| G31_0 | 44 | 32 | 93, 99, 127, 131, 132, 162, 185, 229, 232, 257, 259, 260, 262, 284, 290, 292, 293, 298, 299, 365, 370, 371, 385, 386, 387, 391, 393, 400, 401, 408, 430, 448 | m54118_190409_113909/33685705/ccs | 79 |
| G31_0 | 55 | 32 | 55, 77, 107, 113, 114, 141, 162, 166, 185, 196, 227, 228, 229, 232, 250, 251, 261, 290, 295, 385, 386, 387, 393, 398, 400, 401, 430, 435, 436, 437, 441, 442 | m54118_190409_113909/12059361/ccs | 76 |
| G31_0 | 169 | 33 | 37, 55, 69, 70, 77, 107, 113, 114, 120, 185, 210, 211, 230, 232, 284, 311, 372, 373, 374, 385, 386, 387, 393, 401, 406, 408, 415, 426, 427, 430, 432, 435, 436 | m54118_190409_113909/24772909/ccs | 88 |
| G31_1 | 240 | 20 | 37, 77, 93, 107, 113, 114, 128, 131, 132, 154, 156, 171, 192, 196, 219, 227, 250, 251, 430, 433 | m54118_190409_113909/57278616/ccs | 82 |
| G31_2 | 277 | 24 | 93, 148, 161, 175, 189, 193, 196, 206, 228, 232, 233, 234, 239, 242, 244, 246, 255, 258, 274, 393, 396, 400, 407, 430 | m54118_190409_113909/69992684/ccs | 79 |
| G31_0 | 282 | 41 | 41, 42, 55, 69, 83, 148, 161, 185, 230, 232, 239, 242, 244, 246, 284, 290, 293, 311, 356, 360, 361, 362, 363, 365, 367, 372, 373, 374, 385, 386, 387, 391, 393, 400, 401, 408, 411, 421, 430, 444, 445 | m54118_190409_113909/4588514/ccs | 86 |
| G31_1 | 391 | 12 | 41, 42, 88, 120, 127, 141, 154, 166, 188, 228, 247, 269 | m54118_190409_113909/7013352/ccs | 93 |
| G31_1 | 396 | 104 | 69, 70, 82, 83, 85, 87, 92, 94, 96, 101, 107, 120, 121, 122, 125, 131, 134, 137, 153, 159, 162, 205, 207, 209, 210, 215, 219, 223, 224, 227, 228, 229, 230, 233, 234, 235, 238, 239, 242, 250, 251, 262, 265, 267, 269, 272, 274, 275, 276, 278, 279, 280, 283, 293, 295, 310, 311, 312, 314, 315, 316, 319, 321, 325, 338, 339, 343, 344, 348, 349, 350, 351, 352, 353, 356, 360, 362, 365, 373, 385, 386, 387, 389, 391, 393, 394, 400, 404, 408, 416, 429, 430, 434, 435, 442, 444, 450, 452, 453, 454, 455, 457, 460, 461 | m54118_190409_113909/7668711/ccs | 87 |
| G31_0 | 455 | 16 | 120, 141, 147, 176, 185, 199, 200, 201, 230, 232, 239, 251, 257, 265, 295, 430 | m54118_190409_113909/55705866/ccs | 93 |

Table 3: Summary of grouped CCS reads aligned to G31 haplotypes recovered by `Gretel`. CCS reads (with at least 20 subreads) aligned to the same `Gretel` haplotype with the same combination of mismatch positions are considered identical and counted (#CCS). The CCS read with the highest number of subreads (#Subreads) is chosen as the representative read for that combination of mismatches. The table enumerates mismatch combinations with at least 10 supporting identical CCS reads. Mismatches range from 12 to 104. The SNP positions are on the same co-ordinate system as the graphics in the following section. Care must be taken when interpreting the number of CCS reads that are grouped together (#CCS) as this is primarily influenced by the process of PCR, not by some underlying biological metric such as abundance in the original sample.

#### 11.6 Illumina data do not support G31 CCS reads

##### 11.6.1 Illumina alignment proportions

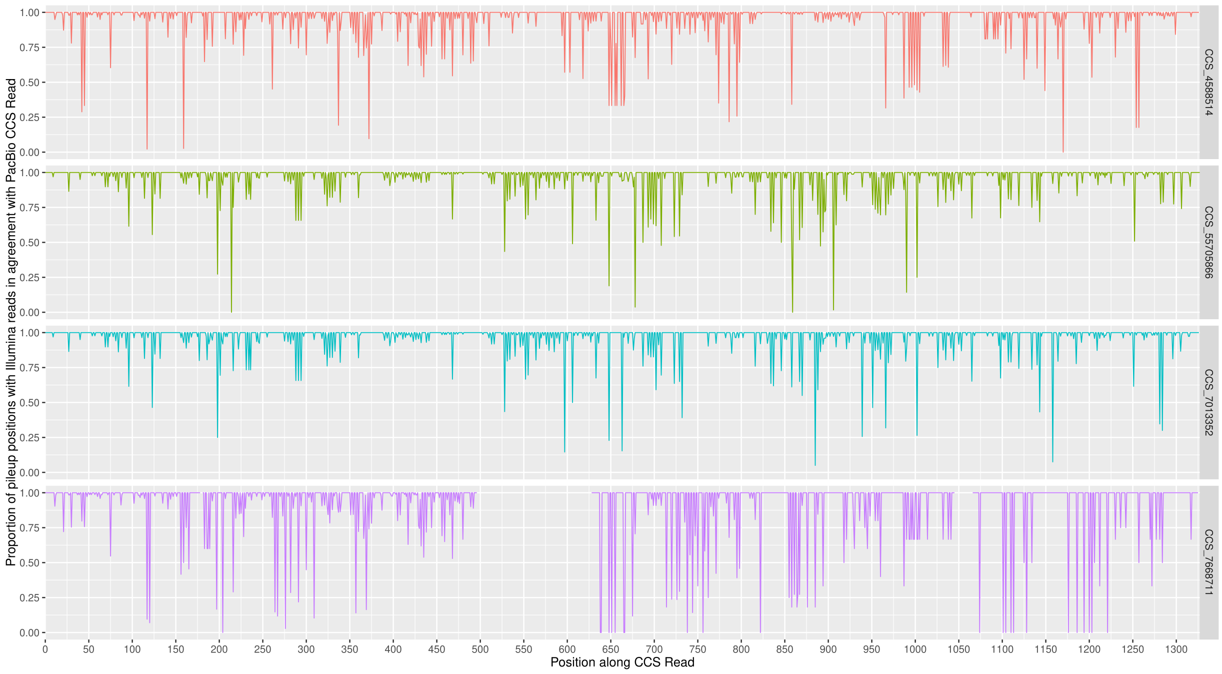

Figure 2: Illumina reads from the initial metatranscriptomic study were aligned to the four most common G31 CCS groups (Table 3) with `minimap2` (`-x sr --end-bonus=8 --secondary=no`). For each position along the CCS read (x-axis), the proportion of aligned Illumina reads that agreed with the CCS read was calculated (y-axes) using `pysam`. The line plots highlight a number of bases for each of the four chosen representative CCS reads (row facets) which have weak support from the corresponding Illumina data. That is, there are positions in the PacBio CCS reads that the Illumina reads do not support.

##### 11.6.2 Illumina alignment pileup examples

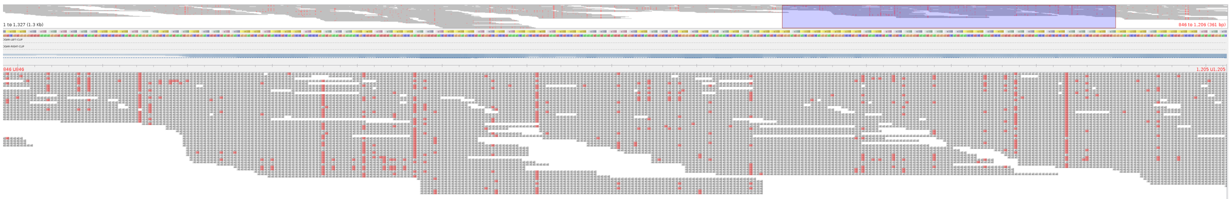

Figure 3: `Tablet` screenshot of Illumina reads aligned against m54118_190409_113909/7013352/ccs with 12 mismatches to the closest `Gretel` haplotype for G31 (Table 3). Red cells indicate bases on Illumina reads that disagree with the PacBio CCS read, highlighting bases with potentially insufficient evidence in the original Illumina sample to have recovered the variants observed on the CCS read.

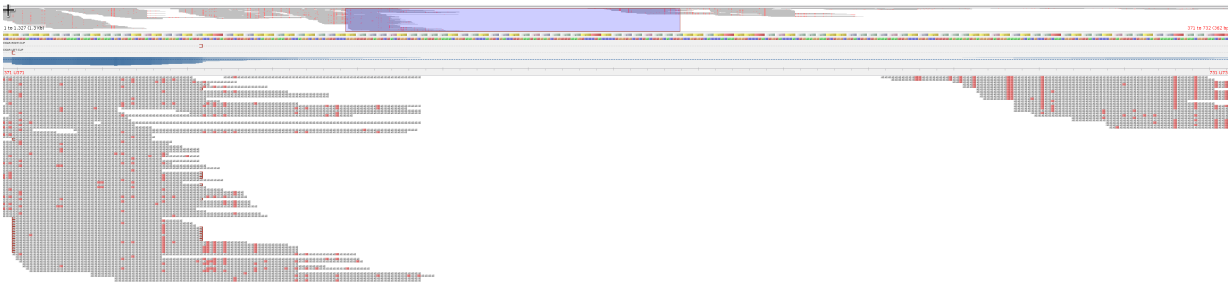

Figure 4: `Tablet` screenshot of Illumina reads aligned against m54118_190409_113909/7668711/ccs with 104 mismatches to the closest `Gretel` haplotype for G31 (Table 3). Red cells indicate bases on Illumina reads that disagree with the PacBio CCS read. The pileup features a large gap, indicating a region on the PacBio read that was not observed by a single Illumina read in the original sample.

##### 11.6.3 Hansel matrix observations

This section contains a set of binary heatmaps, depicting whether more than one observation exists in the `Hansel` matrix to support the transition between two SNPs along a `Gretel` haplotype (upper-diagonal) or a particular PacBio CCS read (lower-diagonal). The `Hansel` matrix stores the pairwise SNP observations parsed from the alignment of short Illumina reads against the reference loci for G31. Dark tiles indicate one or fewer observations support a transition between two SNPs (for any xy co-ordinate in the heatmap). Note the strips of dark tiles on the lower-diagonal indicate that SNPs on the sequenced PacBio CCS read are not found in the original Illumina metatranscriptome, precluding `Gretel`'s ability to recover them. The x and y axes represent co-ordinates along the SNPs recovered in the haplotypes (not the positions within the haplotype itself), matching those in Table 3.

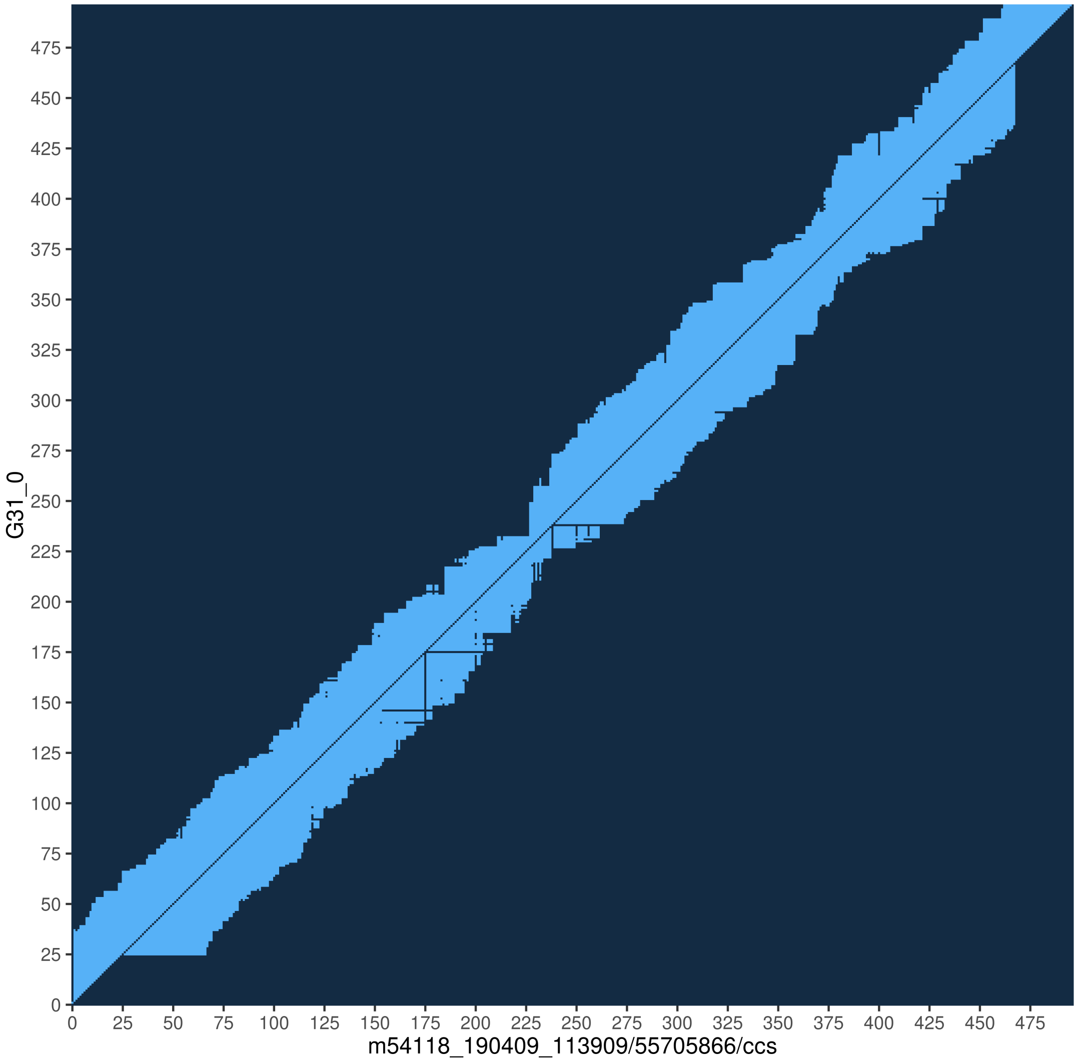

Figure 5: Heatmap depicting SNP pair support for `Gretel` haplotype G31_0 (upper diagonal) and PacBio read m54118_190409_113909/55705866/ccs (lower diagonal) which features 16 mismatches in comparison to G31_0. Light tiles indicate for a given x-y co-ordinate that SNP x and SNP y were observed on at least one read in the Illumina alignment used to generate haplotypes. Conversely, dark tiles indicate one or zero observations of a SNP pair. Note the missing evidence for the PacBio haplotype.

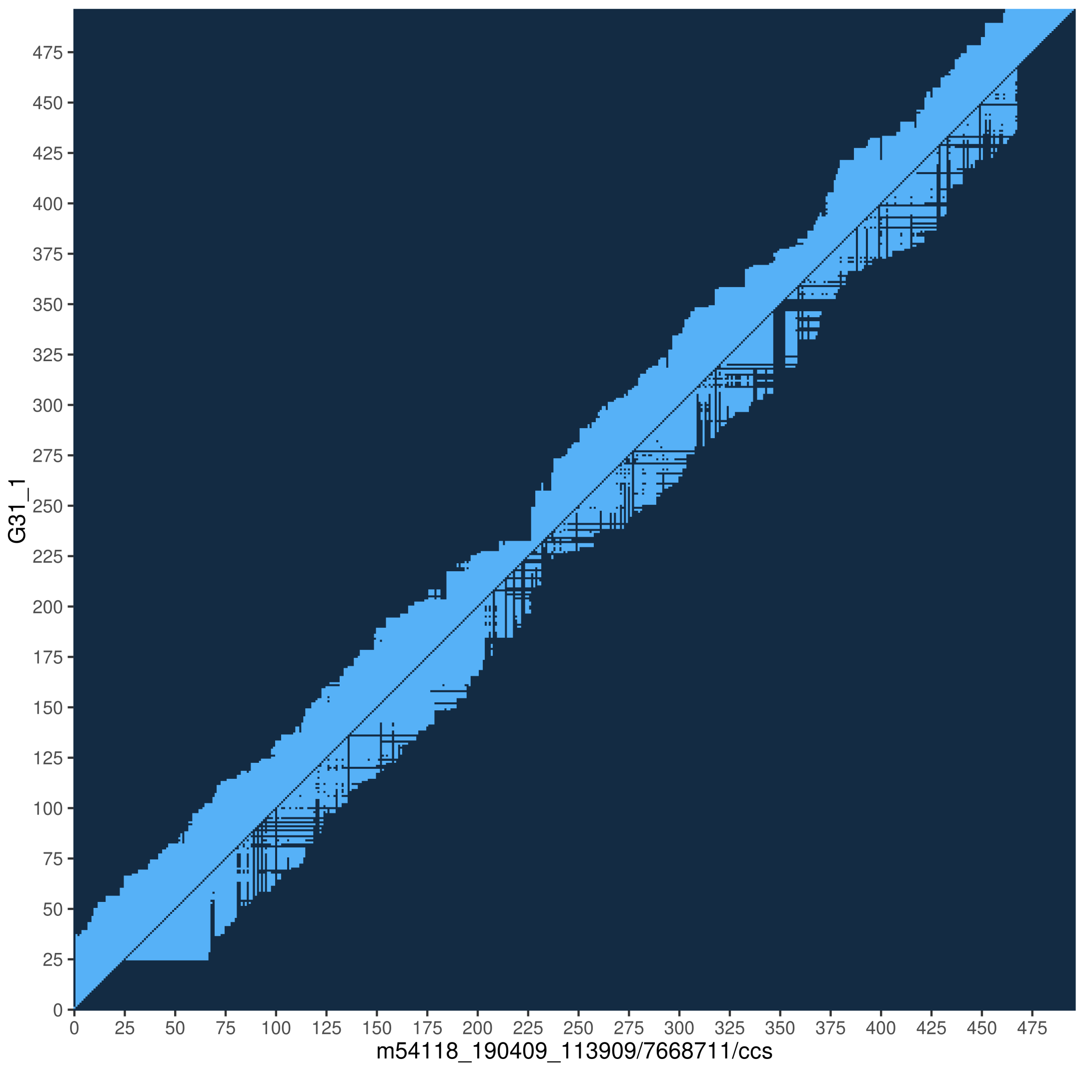

Figure 6: Heatmap depicting SNP pair support for `Gretel` haplotype G31_1 (upper diagonal) and PacBio read m54118_190409_113909/7668711/ccs (lower diagonal) which features 104 mismatches in comparison to G31_1. Light tiles indicate for a given x-y co-ordinate that SNP x and SNP y were observed on at least one read in the Illumina alignment used to generate haplotypes. Conversely, dark tiles indicate one or zero observations of a SNP pair. Note the missing evidence for the PacBio haplotype.

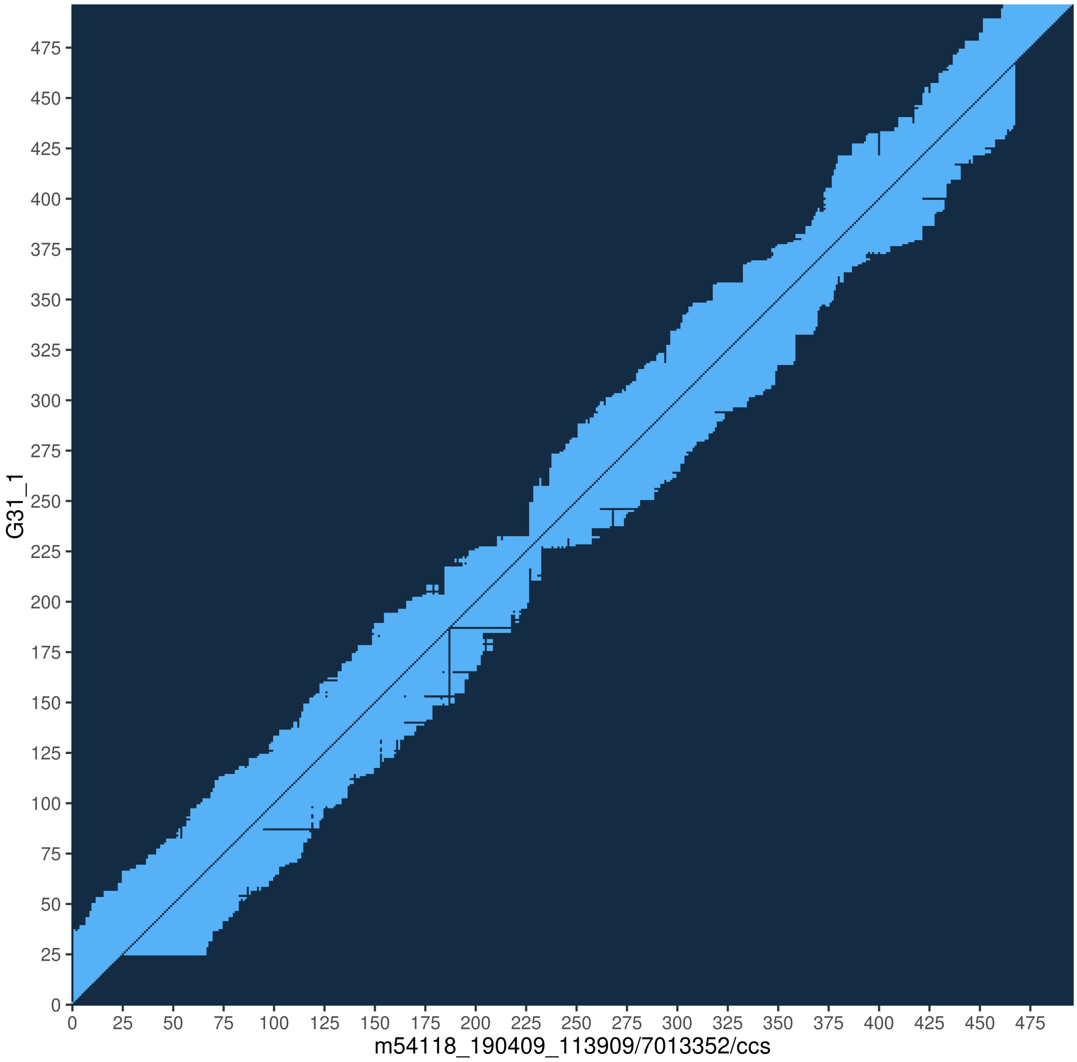

Figure 7: Heatmap depicting SNP pair support for `Gretel` haplotype G31_1 (upper diagonal) and PacBio read m54118_190409_113909/7013352/ccs (lower diagonal) which features 12 mismatches in comparison to G31_1. Light tiles indicate for a given x-y co-ordinate that SNP x and SNP y were observed on at least one read in the Illumina alignment used to generate haplotypes. Conversely, dark tiles indicate one or zero observations of a SNP pair. Note the missing evidence for the PacBio haplotype.

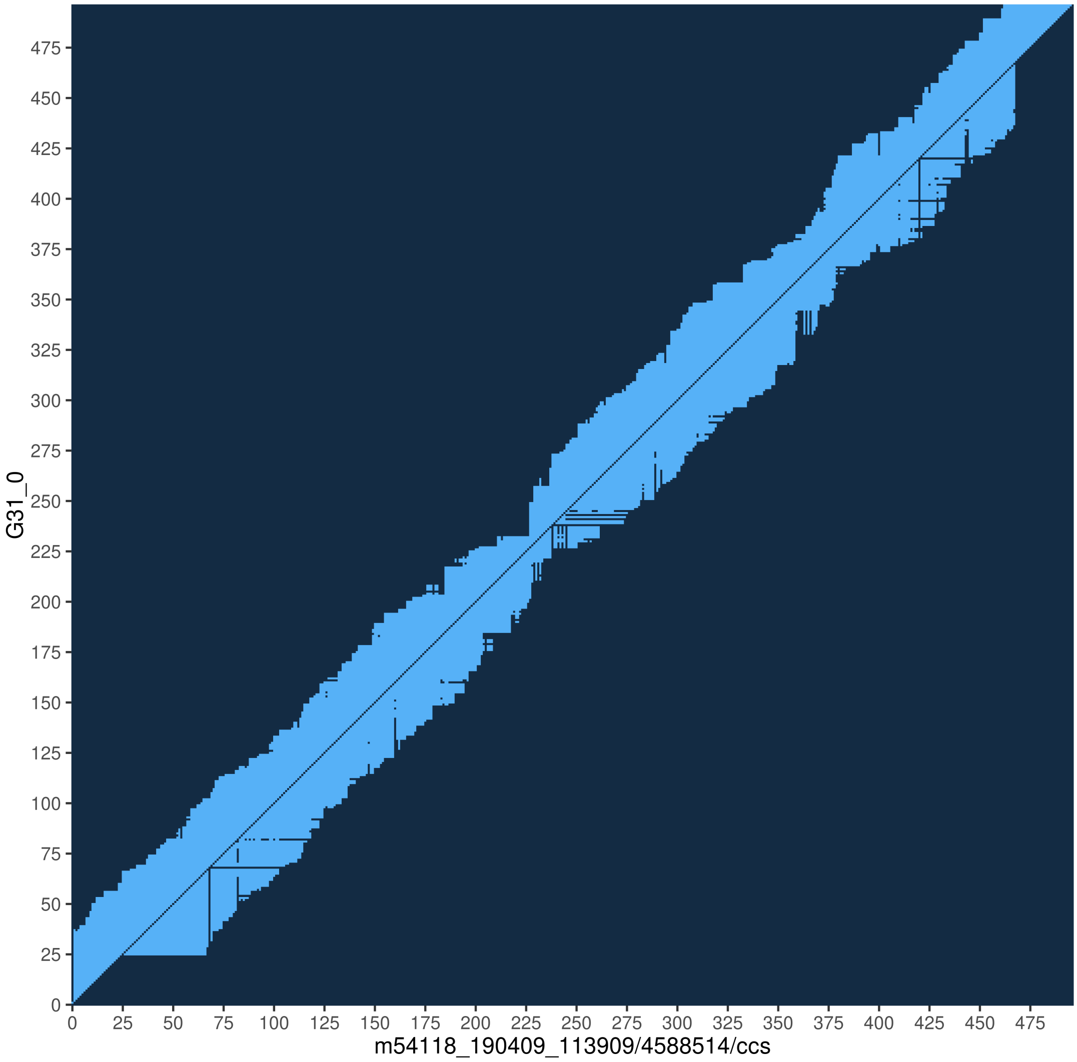

Figure 8: Heatmap depicting SNP pair support for `Gretel` haplotype G31_0 (upper diagonal) and PacBio read m54118_190409_113909/4588514/ccs (lower diagonal) which features 41 mismatches in comparison to G31_0. Light tiles indicate for a given x-y co-ordinate that SNP x and SNP y were observed on at least one read in the Illumina alignment used to generate haplotypes. Conversely, dark tiles indicate one or zero observations of a SNP pair. Note the missing evidence for the PacBio haplotype.

### 12 Hansel and Gretel reveals the haplotype landscape of a pangenome

| Species | Strain | Reads | Regions | Haplotypes† |
|---|---|---|---|---|
| **Ruminococcus flavefaciens** | *007c* | 2,387,278 | 1099 | 7184 |
| | *17* | 1,765,915 | 1054 | 6814 |
| | *AE3010* | 2,520,046 | 1141 | 7597 |
| | *ATCC_19208* | 456,132 | 550 | 3930 |
| | *FD-1* | 3,129,452 | 2178 | 15965 |
| | *MA2007* | 2,041,666 | 1056 | 7333 |
| | *MC2020* | 1,873,628 | 887 | 6105 |
| | *ND2009* | 2,552,008 | 1508 | 11945 |
| | *SAb67* | 2,814,161 | 1950 | 13886 |
| | *XPD3002* | 1,570,131 | 1144 | 8807 |
| | *Y1* | 2,107,083 | 1055 | 7333 |
| | *YAD2003* | 2,424,753 | 1072 | 7093 |
| | *YL228* | 1,785,929 | 1087 | 7827 |
| | *YRD2003* | 2,048,576 | 1020 | 7332 |

Table 4: The 14 Hungate references used for the rumen landscape study. Read sets from the two chosen samples were individually aligned to each of the reference sequences. For each reference, this table reports the total number of reads aligned, the number of regions that returned more than one haplotype (for dN/dS calculations) and the total number of haplotypes recovered from these regions († including duplicate haplotypes).

### 13 Metahaplomes from real reads: HIV 5 strain mix

#### 13.1 Methods

For verification of our approach on real sequenced reads, we evaluated `Gretel` with a dataset designed specifically for the benchmarking of haplotype reconstruction methods [1]. Five well studied HIV-1 strains (*89.6*, *HXB2*, *JRCSF*, *NL43*, and *YU2*) were mixed and sequenced with an Illumina MiSeq.

The Illumina reads are available from ENA (*SRR961514*). As per our protocol, reads were aligned against a pseudo-reference. We selected one of the five strains to serve as this reference (*89.6*), and aligned all sequenced reads against it using `bowtie2`. The overall alignment rate was 96.87%, yielding an alignment of 1,385,162 sequences.

We determined that any heterozygous pileup in the resulting alignment would be defined as a SNP, resulting in a VCF containing 9,570 called variants. The SNPs are so numerous that they occur at 98.98% of all sites.

We executed `Gretel` on five of the largest genes on the HIV-1 genome, using *HXB2* gene co-ordinates [2], each time providing the same alignment BAM and VCF to `Gretel`, but additionally using the `start` and `end` command line parameters to define the boundary of the particular region of interest. Table 5 describes the five genes and their properties. Table 6 shows the identity matrix for the five strains.

We evaluated our approach using the same framework as our synthetic metahaplomes. We report the proportion of variants correctly recovered by `Gretel` for each of the five known haplotypes, for each of the genes.

#### 13.2 Results

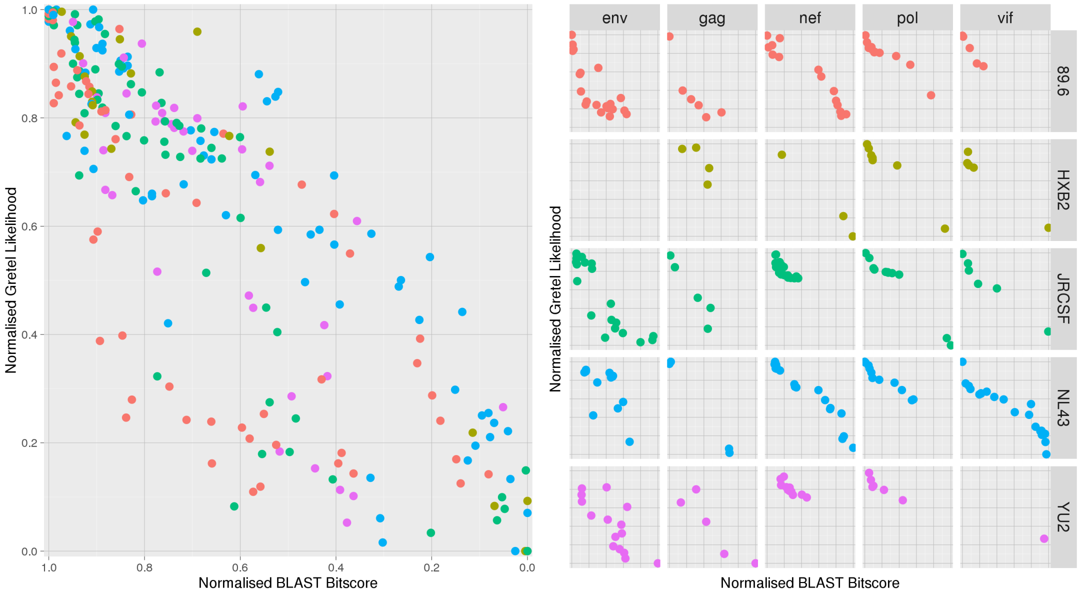

Figure 9: Scaled `BLAST` bitscores (x-axes) against scaled `Gretel` likelihoods (y-axes). (Left) Bitscores against likelihoods for all recovered HIV-1 gene haplotypes, coloured by strain, (Right) Plot facets show the bitscores against likelihood for all recovered haplotypes, separated by gene (column facets) and strain (row facets and colours). `Gretel` consistently awards higher likelihoods to recovered haplotypes that are better matches to a real haplotype.

Finally we apply our approach to a set of real reads (ENA:*SRR961514*) consisting of five distinct HIV strains mixed *in vitro* and sequenced on an Illumina MiSeq [1]. Although the sequences of the five strains were known prior to mixing, the strains are likely to have mutated prior to the sequencing of the samples, introducing cryptic diversity [1]. Furthermore, the sequencing process can add additional noise through sequencing error. This provides a challenging data set for haplotype recovery. For the five longest genes on the HIV-1 genome (gag, pol, vif, env and nef) we used `Gretel` to recover all haplotypes present using the real sequencing reads from this experiment.

Table 7 shows, for each gene from each original HIV strain, the similarity of the nearest matching reconstructed haplotype. We found that the *env* gene had the most novel diversity, with the closest matching haplotypes to the original HIV strains only occurring in the top 25 most likely reconstructions. This is in contrast to *pol*, where the closest matching haplotypes to the original HIV strains were in the top 6 most likely reconstructions. This likely represents differing numbers of novel cryptic haplotypes of these genes and correlates well with their known mutation rates *in vivo* [3].

For all of the recovered HIV-1 haplotypes, Figure 9 plots the scaled `BLAST` bitscores against the `Gretel` likelihood score. Each recovered haplotype is matched to its closest strain. Our results show that ordering the recovered haplotypes by their likelihood scores can be used as a method to find the best recoveries amongst the recovered sequences.

As reported in Table 7, haplotypes were discarded if they did not meet a conservative threshold of $-1000\ log_{10}$ likelihood. Manual inspection indicated that these discarded haplotypes showed over selection of deletions, an artifact arising from `Gretel` exhausting non-deleterious evidence in `Hansel` before terminating. Such haplotypes yield no significant BLAST hits against the NCBI nr database [4] and our likelihoods provided a clear distinction between noise and useful recoveries.

Beyond the alignment, Gretel does not require read processing, parameter bootstrapping or error correction. Despite thousands of heterogeneous sites (primarily caused by sequencing error and alignment noise) Gretel is still capable of recovering known haplotypes with 100% accuracy and biologically relevant cryptic haplotypes from metagenomic data.

| **Gene** | **Region (Size)** | **Average Coverage** |
|---|---|---|
| *gag* | 790—2289 (1,500 bp) | 32,325.34 |
| *pol* | 2084—5093 (3,009 bp) | 46,984.47 |
| *vif* | 5040—5616 (576 bp) | 24,110.89 |
| *nef* | 8796—9414 (618 bp) | 21,059.34 |
| *env* | 6224—8792 (2,568 bp) | 22,699.35 |

Table 5: The five HIV-1 genes, HXB2 co-ordinates [2] and associated properties of the aligned reads

| **Strain** | *89.6* | *JRCSF* | *YU2* | *HXB2* | *NL43* |
|---|---|---|---|---|---|
| *89.6* | 100.00 | – | – | – | – |
| *JRCSF* | 93.98 | 100.00 | – | – | – |
| *YU2* | 94.05 | 95.10 | 100.00 | – | – |
| *HXB2* | 94.43 | 95.18 | 95.71 | 100.00 | – |
| *NL43* | 94.07 | 94.98 | 95.39 | 97.49 | 100.00 |

Table 6: Percentage identity matrix for the five HIV-1 strains as reported by MUSCLE [5]

| | | | **Strains** | | | | |
|---|---|---|---|---|---|---|---|
| **Gene** | **SNPs** | **Haplotypes†** | **89.6** | **HXB2** | **JRCSF** | **NL43** | **YU2** |
| *gag* | 1500 | 24 | **100.0** (2) | **100.0** (6) | 99.33 (4) | **100.0** (3) | 99.40 (10) |
| *pol* | 3009 | 38 | 99.73 (4) | 99.20 (3) | 99.67 (2) | **100.0** (1) | 98.44 (6) |
| *vif* | 576 | 38 | **100.0** (2) | 98.96 (9) | **100.0** (3) | **100.0** (1) | 97.40 (5) |
| *nef* | 618 | 60 | **100.0** (2) | 97.14 (6) | 97.30 (15) | 97.62 (5) | 96.02 (12) |
| *env* | 2568 | 66 | 99.96* (1) | 97.90 (11) | 99.69 (7) | 98.83 (12) | 99.45 (25) |

Table 7: For each gene from each original HIV strain, the percentage similarity of the nearest matching reconstructed haplotype. Results are presented for each of the five longest genes of the HIV-1 genome (*gag*, *pol*, *vif*, *nef*, *env*). The bracketed figure indicates the rank of the best haplotype for the strain amongst all recovered haplotypes, according to its likelihood score. We also report the total number of haplotypes recovered by Gretel for each gene. *Recovery of 89.6 *env* gene has just one incorrect SNP (2567 SNPs recovered). †Number of haplotypes returned after conservative -1000 $log_{10}$ likelihood cutoff.

### References

[1] Francesca Di Giallonardo, Armin Töpfer, Melanie Rey, Sandhya Prabhakaran, Yannick Duport, Christine Leemann, Stefan Schmutz, Nottania K. Campbell, Beda Joos, Maria Rita Lecca, Andrea Patrignani, Martin Däumer, Christian Beisel, Peter Rusert, Alexandra Trkola, Huldrych F. Günthard, Volker Roth, Niko Beerenwinkel, and Karin J. Metzner. Full-length haplotype reconstruction to infer the structure of heterogeneous virus populations. *Nucleic Acids Res*, 42(14):e115, 2014.

[2] Bette Korber, Brian T Foley, C Kuiken, Satish K Pillai, Joseph G Sodroski, et al. Numbering positions in HIV relative to HXB2CG. *Human retroviruses and AIDS*, 3:102–111, 1998.

[3] J.M. Cuevas, R. Geller, R. Garijo, J. López-Aldeguer, and R. Sanjuán. Extremely high mutation rate of HIV-1 in vivo. *PLoS Biol*, 13(9):e1002251, 2015.

[4] NCBI Resource Coordinators. Database resources of the National Center for Biotechnology Information. *Nucleic Acids Research*, 44(Database issue):D7–D19, 2016.

[5] R. C. Edgar. MUSCLE: a multiple sequence alignment method with reduced time and space complexity. *BMC Bioinformatics*, 5(1):113, 2004.
